# Bacteria combine polar- and dispersed-growth to power cell elongation and wall width dynamics

**DOI:** 10.1101/2024.07.30.605496

**Authors:** Matthew P. Zambri, Christine R. Baglio, Oihane Irazoki, Stephanie E. Jones, Ethan C. Garner, Felipe Cava, Marie A. Elliot

## Abstract

The cell wall is a complex structure. For most bacteria, peptidoglycan is an essential component of their cell wall, with different bacteria having evolved distinct biosynthetic strategies. The mechanisms driving bacterial growth can be divided into three, mutually-exclusive categories: (i) dispersed growth, mediated by MreB and employed by many rod-shaped bacteria; (ii) polar growth, driven by distinct proteins in the actinobacteria and rhizobiales; and (iii) septal growth, fueled by FtsZ in many coccoid bacteria. Here, we show that under conditions of rapid growth, the actinobacterial representative *Streptomyces venezuelae* transcends these categories, simultaneously employing both canonical polar growth, and MreB-mediated dispersed growth. Our results indicate that MreB is essential for cell wall integrity and culture viability under these growth conditions, promotes dynamic cell wall changes over the course of a growth cycle, and contributes to a wall that is structurally distinct from that of conventionally growing streptomycetes.

## Introduction

All organisms face the challenge of coupling cell growth and expansion with cell wall synthesis, and the need to adjust this process in response to changing environmental conditions. How this challenge is managed depends on the organism, as cell wall growth and its control can be achieved by diverse mechanisms. Polar (tip) growth is one strategy employed across kingdoms, driving the expansion of everything from animal neurites and plant root hairs, to filamentous fungi and select bacteria (*e.g.*, the actinobacteria and rhizobiales).

Within the polar-growing actinobacteria, *Streptomyces* are one of the best-studied groups. These filamentous microbes have a complex life cycle with different developmental trajectories, including the well-established classical sporulating life cycle (Zambri *et al*., 2022), and the more recently described exploratory growth mode (Jones *et al.,* 2017; Jones & Elliot, 2017; Jones & Elliot, 2018). Their multi-staged classical life cycle begins with spore germination and hyphal outgrowth, and the ensuing vegetative growth involves hyphal tip extension and branching. The subsequent reproductive growth phase involves raising polarly-growing but non-branching aerial hyphae, which are subsequently converted into chains of dormant spores. Exploratory growth also begins with spore germination. This is followed by a rapid expansion of vegetative-like hyphal filaments across solid growth surfaces, at a rate that is an order of magnitude faster than that observed during classical growth. The mechanistic basis by which *S. venezuelae* achieves such rapid growth has been a major question in the field.

Cell growth is mediated in part by cell wall expansion, with peptidoglycan being an integral component of the bacterial cell wall. In *Streptomyces,* DivIVA is the key driver of growth at the hyphal poles (Flӓrdh, 2003). DivIVA is broadly conserved among Gram-positive bacteria (Hammond, White & Eswara, 2019), but its role in cell wall synthesis is confined to the actinobacteria, where it is an essential protein (Flӓrdh, 2003). In *Streptomyces,* DivIVA localizes to the tips of vegetative and aerial hyphae as part of the ‘polarisome’, where it recruits the peptidoglycan biosynthetic machinery.

In contrast to the DivIVA-mediated polar growth of *Streptomyces*, many rod-shaped bacteria (*e.g., Escherichia coli* and *Bacillus subtilis*) instead insert new peptidoglycan along their lateral side walls. This dispersed wall growth is governed by MreB, which recruits and organizes the peptidoglycan biosynthetic proteins into an ‘elongasome’ or ‘rod’ complex (Salje *et al.,* 2011; Muchová, Chromiková & Barák, 2013; Shi *et al.,* 2018). Along the length of the cell, MreB assembles into discrete filaments that move circumferentially within the cell as new peptidoglycan is inserted (Jones, Carballido-López & Errington, 2001; van den Ent *et al.,* 2010; Garner *et al.,* 2011; van Teeffelen *et al.,* 2011; van den Ent *et al.,* 2014). *Streptomyces* have traditionally been considered unusual amongst the polarly-growing bacteria in encoding both polar and lateral wall growth determinants, with DivIVA driving active polar growth, and MreB activity being confined to spore wall maturation during classical reproductive growth (Mazza *et al.,* 2006; Heichlinger *et al.,* 2011).

Historically, the distinction between bacteria growing via polar wall insertion versus those employing dispersed wall synthesis has been absolute. Here, we report that *S. venezuelae* employs polar and dispersed cell wall synthesis during exploratory growth. This hybrid growth mode requires both DivIVA and MreB to build a uniquely dynamic peptidoglycan structure with altered compositional properties.

## Results

### *S. venezuelae* alters its cell wall structure and composition during exploration

Exploring *S. venezuelae* colonies expand at a rate that is >10× faster than classically growing vegetative colonies (Jones *et al.,* 2017), although the onset of this rapid expansion can vary depending on growth conditions. Exploration can be readily induced by growth on yeast extract and peptone (YP) (**Video S1**), or by growth on YP with D-glucose in the presence of the yeast *Saccharomyces cerevisiae* (a condition termed YPDy) (Jones *et al*., 2017). The exploring phenotypes on YP and YPDy media are similar, although the process unfurls more quickly on YP; growth on YPDy is associated with an initial delay in the initiation of rapid exploratory growth (Jones *et al.,* 2017).

To determine if the enhanced colony expansion associated with exploratory growth involved changes to the cell wall of exploring hyphal filaments, we first employed transmission electron microscopy (TEM). We examined wild type cultures grown on YP medium over an 11-day time course and measured the cell wall thickness of exploring hyphae at 2-day intervals, starting at day 5. We focused our attention on hyphae at the leading edge of the colonies, as this was expected to be the site of most active growth. Intriguingly, for the earliest explorers examined (5 days), we observed a significant thinning (∼20%) of the cell wall relative to classically growing vegetative hyphae. As exploration proceeded, we observed a stepwise increase in cell wall thickness, culminating in a median wall thickness at day 11 that exceeded that of the day 5 hyphae by >40% (**Fig. 1A**). Similar trends were observed for YPDy-grown cultures (**Fig. S1A**). Interestingly, cell wall thickening during exploratory growth did not correlate with increased hyphal width (**Fig. S2**).

**Figure 1:**
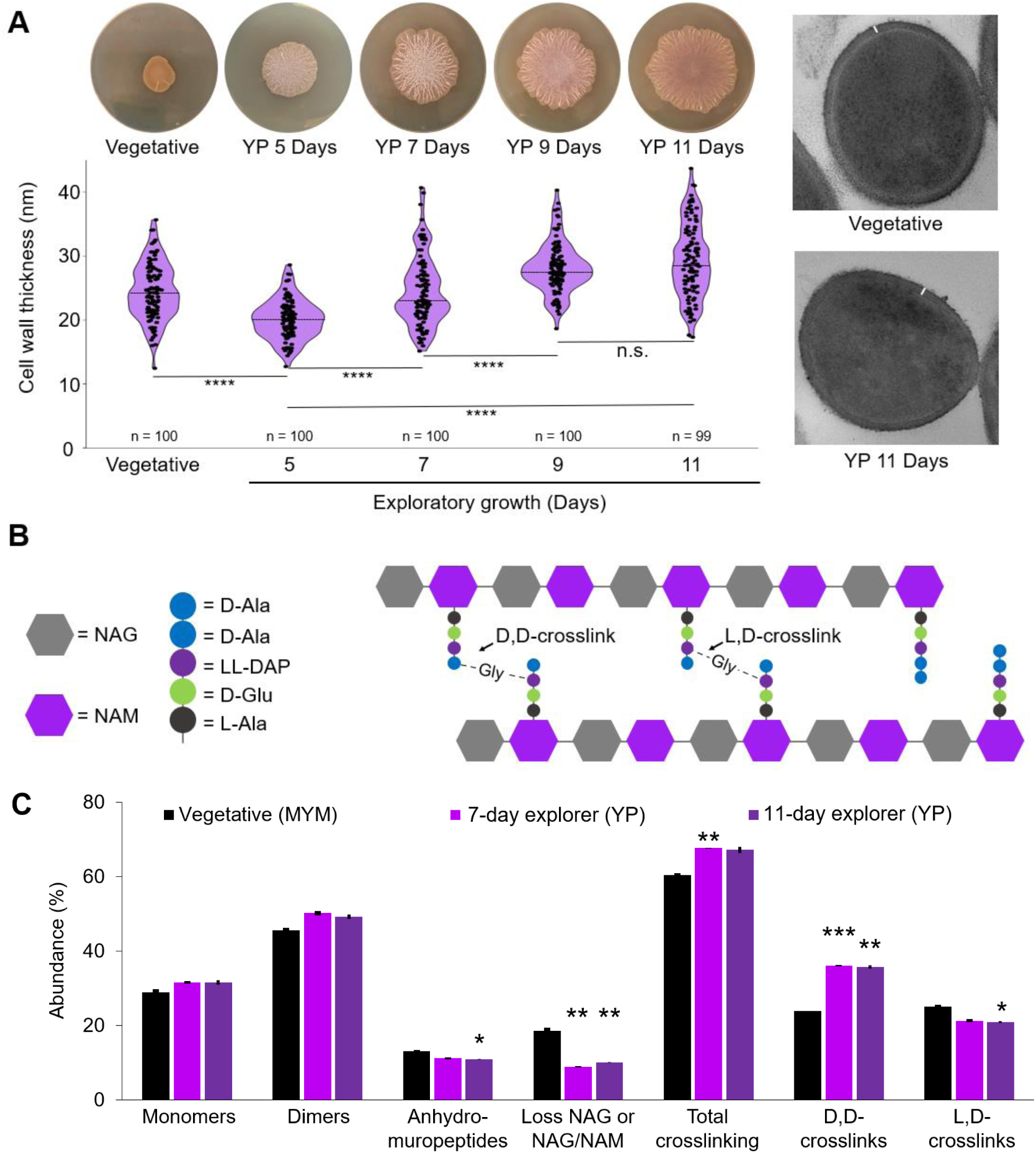
Exploring cell walls differ from those of classically-growing vegetative cells in *S. venezuelae*. **(A)** Left: Violin plot of hyphal cell wall thickness for vegetative cells (MYM-grown) and exploring cells (YP-grown) at different stages in the exploration cycle, as determined by measuring transmission electron micrographs of each using ImageJ. Three measurements were taken from distinct positions around each hypha, and averaged. Above each plot are representative images of vegetative/exploring colonies from the time points indicated. Right: representative TEM micrographs of vegetative (top) and exploring hyphae (bottom), with the white lines indicating a cell wall width measurement point. Statistical significance was determined via Mann-Whitney tests (**** = P value < 0.0001). **(B)** Peptidoglycan schematic, where it is composed of alternating subunits of *N*-acetylglucosamine (NAG) and *N*-acetylmuramic acid (NAM) connected via β-(1,4) glycosidic linkages. These strands bear peptide stems that can be attached through either D,D-or L,D-crosslinks. The peptide stem of *Streptomyces* is typically L-Ala—D-Glu—LL-DAP—D-Ala—D-Ala. **(C)** Bar graph comparing peptidoglycan analyses of wild type vegetative *S. venezuelae* cells (MYM-grown) and explorer cells (YP-grown) after 7-and 11-days. Shown are the abundances of total muropeptide monomers, dimers, anhydromuropeptides, muropeptides lacking NAG or NAG/NAM, total crosslinking, total D,D-crosslinks and total L,D-crosslinks, following peptidoglycan isolation, digestion and UPLC-MS analysis. Asterisks indicate statistically significant differences between vegetative samples and exploring samples (* = P value < 0.05, ** = P value < 0.01, *** = P value < 0.001, **** = P value < 0.0001; as determined by Student’s t-tests). Error bars indicate standard error. Data presented are the average values of either three replicates (7-and 11-day YP-grown explorers) or two replicates (MYM-grown vegetative cells).

Given the dynamic changes in cell wall thickness observed over an exploration cycle, we wondered whether the composition or structure of cell walls from exploring colonies might also differ from those of classically growing vegetative cells, and if this changed over time. The peptidoglycan backbone typically comprises alternating subunits of *N-*acetylglucosamine (NAG) and *N*-acetylmuramic acid (NAM). These glycan strands are joined via their NAM-affixed peptide stems, with these stems in turn being linked together by either D,D-crosslinks that are catalyzed by penicillin-binding proteins (PBPs), or L,D-crosslinks catalyzed by L,D-transpeptidases (Glauner, Höltje & Schwarz, 1988) (**Fig. 1B**). We isolated peptidoglycan from: (i) cultures grown on YP and YPDy agar over the course of an exploration cycle, (ii) vegetative cultures grown on the classical growth medium MYM (malt extract, yeast extract, and maltose), and (iii) cultures grown on YPD (without yeast, where no exploration is observed, and colony growth is exclusively vegetative). We analyzed their composition using ultra-performance liquid chromatography coupled with mass spectrometry (UPLC-MS) and found exploring hyphae had distinctive peptidoglycan profiles relative to both classical vegetatively growing hyphae (**Fig. 1C**; **Tables S1 and S2**) and previously reported profiles for *Streptomyces coelicolor* (van der Aart *et al.,* 2018). In comparing early/mid stage (7-day) and late stage (11-day) YP-grown explorer cells, with vegetative hyphae grown on MYM, we observed increased total crosslinking (∼7%) and D,D-crosslinking (∼12%), but decreased levels of L,D-crosslinking (∼4%) for the exploring cultures relative to the vegetive culture (**Fig. 1B & C**). A near identical trend was observed for YPDy-grown explorers, albeit over a longer timescale (**Fig. S1B**), consistent with the slower initiation of colony expansion observed during growth on YPDy medium (**Video S2**).

### MreB1 and the *mre* operon are critical for exploratory growth

Having established that exploring cells exhibit unusual cell wall dynamics and structural changes compared with classically growing cells, we hypothesized that specific cell wall biosynthetic factors may function to promote the rapid growth of exploring cells. As MreB is a cell wall growth-promoting determinant in many bacterial species and is broadly conserved in the streptomycetes, we wondered whether it may play a role in facilitating exploratory growth. There are two copies of *mreB* in the *S. venezuelae* genome, dubbed *mreB1* (*vnz_11695*) and *mreB2* (*vnz_11715*) (**Fig. 2A**). We generated individual *mreB1* and *mreB2* mutant strains. Both strains looked effectively wild type when growing classically; however, the Δ*mreB1* strain exhibited strong defects in exploration during growth on YP agar, including smaller overall colony size, abnormal wrinkling patterns, and widespread regions of what appeared to be cell lysis (**Fig. 2B**; **Video S3**). Introducing a wild type copy of *mreB1* at a heterologous site in the chromosome restored a more wild type-like wrinkling pattern and alleviated the apparent cell lysis, although colonies remained smaller than wild type (**Fig. 2B**; **Video S4**). In contrast, the Δ*mreB2* strain was smaller than the wild type strain, but had an otherwise similar morphology and did not appear to undergo lysis (**Fig. S3**). An Δ*mreB1/*Δ*mreB2* double mutant effectively phenocopied the *mreB1* mutant (**Fig. S3**). We also generated an *mre* operon deletion strain lacking *mreB1, mreC, mreD, pbp2,* and *rodA* (*vnz_11675-vnz_11695*) (**Fig. 2A**), where the other genes encode additional elongasome members and are known to participate in spore maturation together with MreB (Kleinschnitz *et al.,* 2011). This strain had a distinct exploration phenotype, in that overall colony size was larger than that of the *mreB1* mutant; however, colony-wide lysis was still observed (**Fig. S3; Video S5**). To probe the apparent cell lysis observed for the mutant *mreB1* and *mre* operon colonies during exploratory growth, we introduced a constitutively expressed *mNeonGreen* construct into the chromosome of these strains and into a wild type control strain, and monitored colony fluorescence over 10 days on YP medium. A complete loss of fluorescence was observed in the mutant strains in the regions of the colonies that appeared to be lysing, while the fluorescence of the wild type strain remained well above background levels at all times (**Fig. 2C**; **Fig S3**). Scanning electron micrographs of Δ*mreB1* exploring cells further revealed an abundance of hyphae with abnormal ‘swollen’ morphologies, alongside many apparently lysed (deflated) hyphae, when compared with wild type (**Fig. 2D**). These observations collectively implicated MreB and its associated elongasome components as being essential for exploratory growth under these conditions.

**Figure 2:**
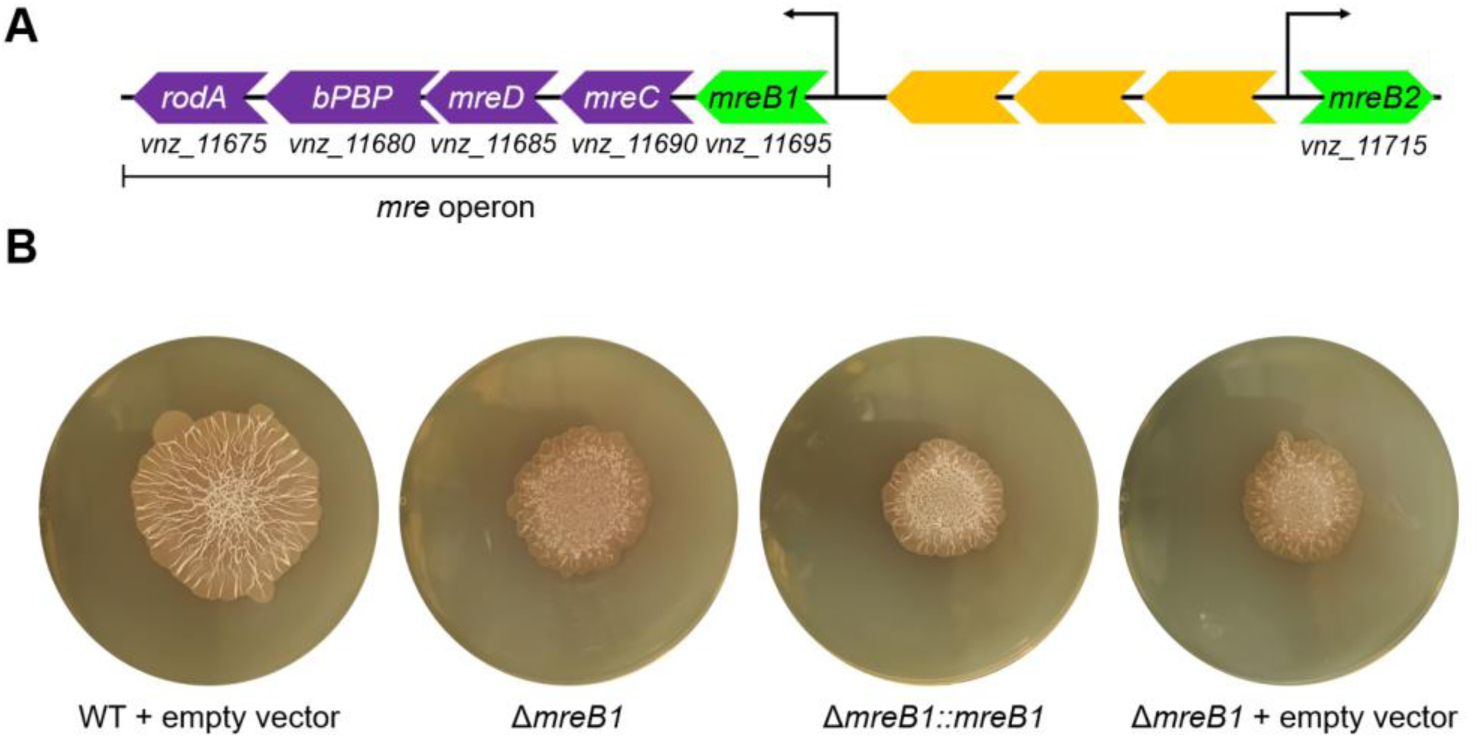
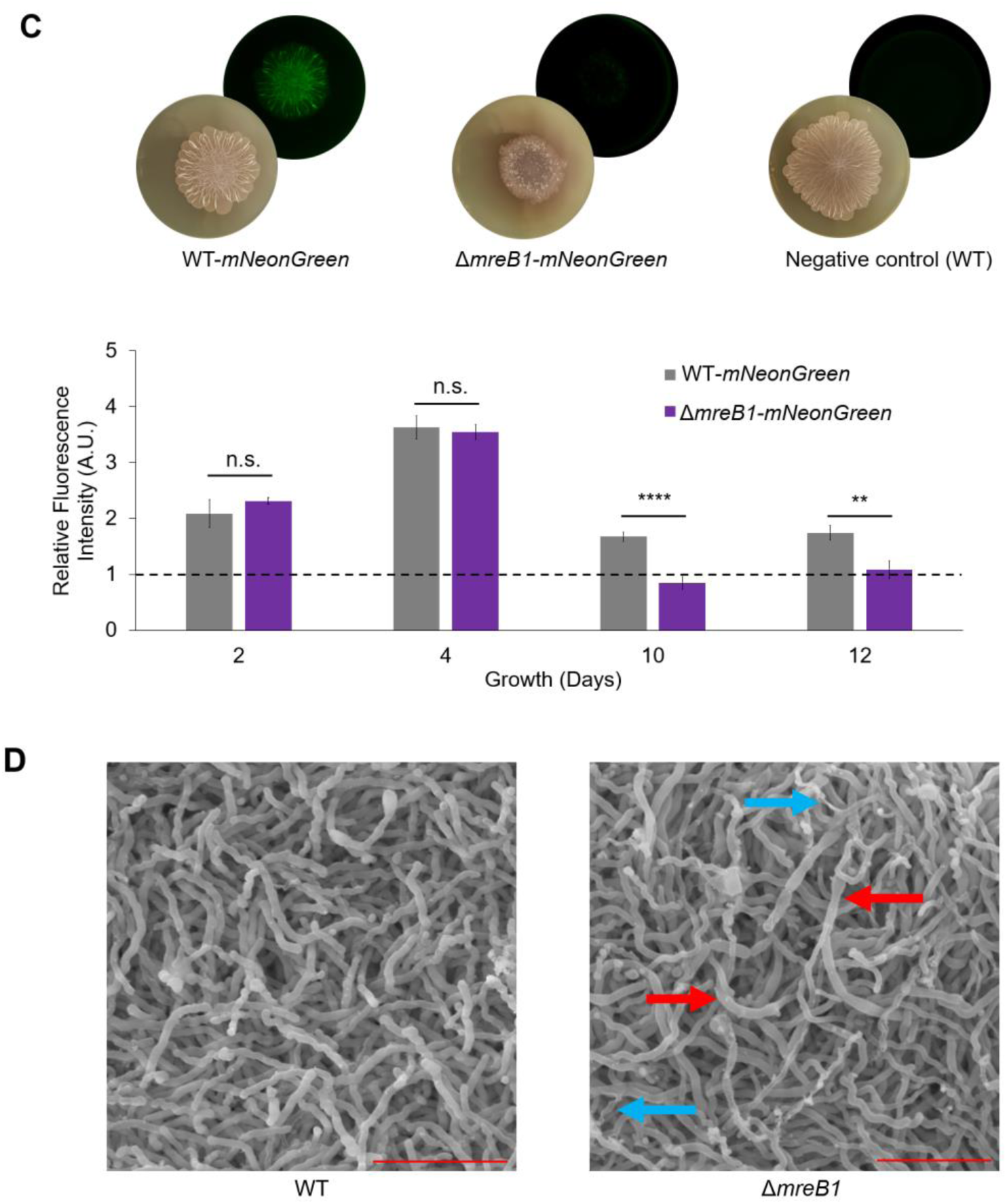
Loss of *mreB1* leads to exploration defects. **(A)** Schematic of the *mre* operon in *S. venezuelae,* which includes *mreB1*, *mreC*, *mreD*, *bPBP*, and *rodA*, with *mreB2* located upstream (*bPBP:* encodes a Class B penicillin-binding protein); their associated transcription start sites are indicated. Schematic is not to scale. **(B)** Wild type (WT) with empty vector, Δ*mreB1*, Δ*mreB1* complemented with *mreB1* on an integrating plasmid vector (Δ*mreB1*::*mreB1*), and Δ*mreB1* carrying the empty vector, grown on YP medium for 11 days. **(C)** Top: Representative light and fluorescence images of exploring colonies grown on YP medium for 10 days. From left to right: WT-*mNeonGreen*, Δ*mreB1*-*mNeonGreen* and a negative control (wild type with empty vector). Bottom: Bar graph comparison of relative fluorescence between wild type and Δ*mreB1* strains with constitutively expressed *mNeonGreen*. Values presented are relative to a negative control strain lacking *mNeonGreen* and are the average of biological triplicates. Cultures were grown on YP exploration medium. Asterisks indicate statistically significant differences (** = P value < 0.01, **** = P value < 0.0001; as determined by Student’s t-tests). Error bars indicate standard error. **(D)** Representative scanning electron micrographs of wild type (WT) and Δ*mreB1* exploring hyphae grown on YP medium for 10 days. Hyphae with abnormal morphologies are highlighted with arrows: red arrows indicate swollen hyphae and blue arrows indicate deflated/lysed hyphae. Red scale bars = 10 μm.

It had been reported previously that supplementing the growth medium with high levels of Mg^2+^ could rescue MreB-deficient *B. subtilis* from cell lysis either by inhibiting peptidoglycan-degrading enzymes (Formstone & Errington, 2005; Tesson *et al.,* 2022; Wilson *et al.,* 2023) or through stabilizing interactions with wall teichoic acids (Kern *et al.,* 2010; Thomas & Rice, 2014). We therefore tested whether Mg^2+^ supplementation could rescue the lysis and/or exploration defects of the *mreB1* and *mre* operon mutant strains. We found that adding 25 mM MgSO_4_ to YP agar led to a moderate exploration defect for the wild type strain; however, it restored wild type-like colony sizes and wrinkling patterns to the mutant strains (**Fig. S4**).

Having established that MreB1 was critical for *S. venezuelae* exploration, we next set out to assess its effect on the cell wall itself. We analyzed the structure and composition of the Δ*mreB1* cell wall during classical growth and exploration on YP medium. TEM micrographs of the Δ*mreB1* strain revealed that the stepwise increase in cell wall thickness observed over the course of an exploration cycle in the wild type strain was not seen for the *mreB1* mutant (**Figs. 1A** and **3A**). Instead, the *mreB1* mutant wall remained significantly thinner than its wild type counterpart at all stages after initiating exploration at 4 days (**Fig. 3B**). The peptidoglycan compositional and structural analyses of the Δ*mreB1* strain differed from that of the wild type, including increased muropeptides lacking sugar moieties, which can be a hallmark of increased peptidoglycan hydrolysis, and decreased total crosslinking, driven by decreases in both D,D-and L,D-crosslinks (**Fig. 3C**; **Tables S1 and S2**). The combination of fewer peptidoglycan crosslinks and possibly increased hydrolysis suggested that the peptidoglycan mesh of the *mreB1* mutant was likely more disordered than that of the wild type strain, and that these cells may be more susceptible to lysis. Importantly, these peptidoglycan changes were only observed in exploring conditions; the peptidoglycan profiles of classically-growing vegetative Δ*mreB1* and wild type strains (MYM-grown) were effectively indistinguishable (**Fig. 3D**). Taken together, these results suggested that MreB is critical for conferring and/or maintaining cell wall integrity in rapidly growing explorer cells.

**Figure 3:**
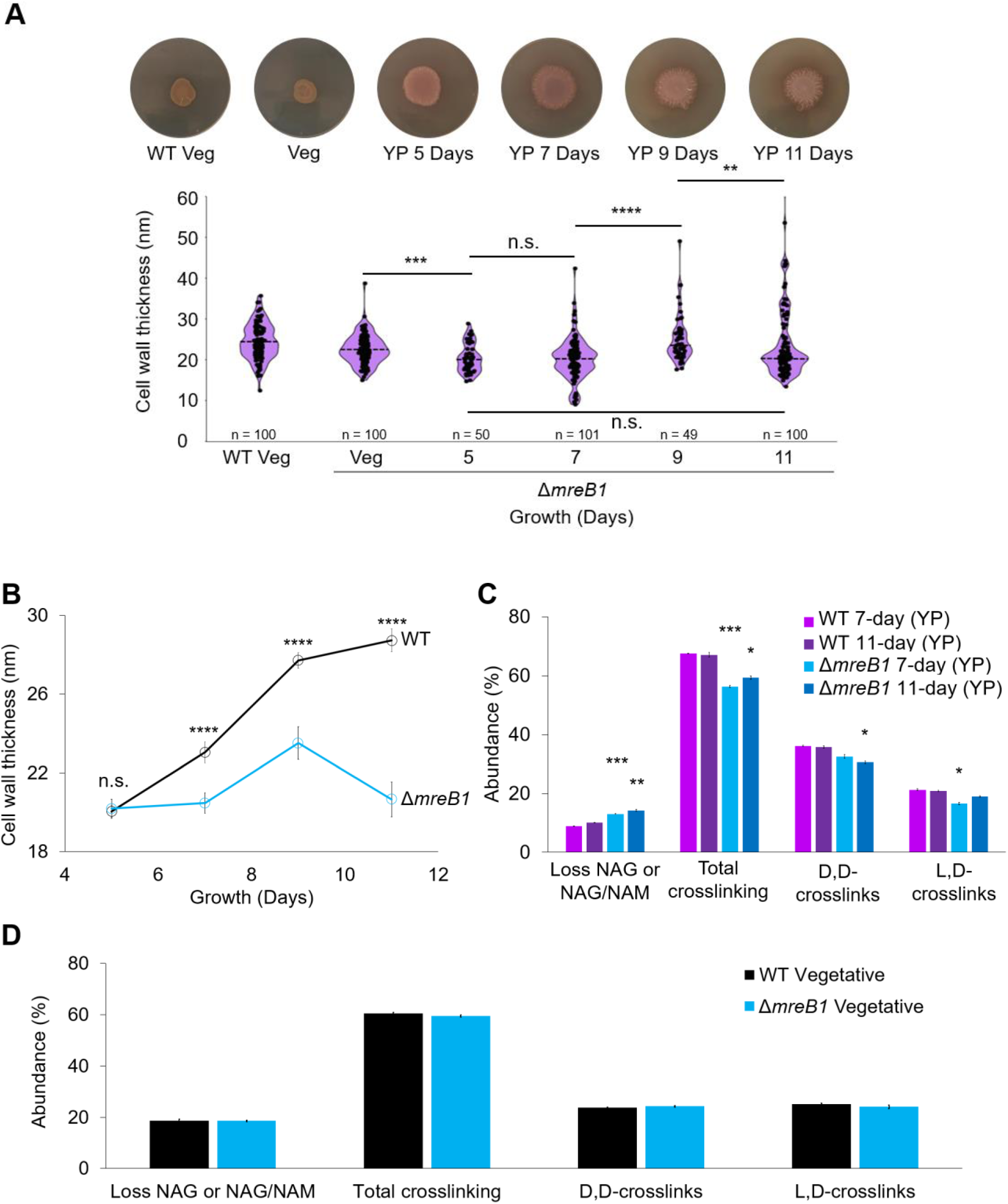
Δ*mreB1* explorers have thinner cell walls and an altered peptidoglycan composition. **(A)** Violin plot of cell wall thickness for intact (non-lysed) hyphae from wild type vegetative (WT Veg; MYM-grown) and Δ*mreB1* vegetative (Veg) and YP-grown exploring cells at different stages of the exploratory cycle (indicated as days post inoculation), as determined by measuring transmission electron micrographs of each using ImageJ. Dashed line = median values. For all hyphae, three measurements were taken per cell and these were averaged before plotting. A Mann-Whitney statistical test was performed to determine statistical significance (n.s. = not significant; *** = P value < 0.001; **** = P value < 0.0001). **(B)** Line plot of median cell wall thickness measurements from exploring wild type and Δ*mreB1* strains. Asterisks indicate statistically significant differences between WT and Δ*mreB1* strains at corresponding time points (*** = P value < 0.001; **** = P value < 0.0001; as determined by Mann-Whitney tests). **(C)** The abundance of total crosslinks, D,D-crosslinks and L,D-crosslinks upon isolation, digestion and UPLC-MS analysis of wild type and Δ*mreB1* peptidoglycan from cultures grown on YP exploration medium. Asterisks indicate statistically significant differences between WT and Δ*mreB1* strains at corresponding time points (* = P value < 0.05, ** = P value < 0.01, *** = P value < 0.001; as determined by Student’s t-tests). Error bars indicate standard error. The data presented are the average values obtained from three biological replicates for each condition. **(D)** The abundance of total crosslinks, D,D-crosslinks and L,D-crosslinks upon isolation and UPLC-MS analysis of digested peptidoglycan of vegetatively growing wild type and Δ*mreB1* samples grown on MYM medium. No statistical differences were observed. Error bars indicate standard error. The data presented are the average values obtained from two biological replicates for each condition.

### Lateral wall synthesis in exploring cultures appears to require MreB1

We next set out to follow cell wall synthesis during exploration, assessing the capacity for DivIVA-driven polar growth, and determining whether MreB-driven dispersed wall synthesis was occurring. To do this, we employed the fluorescent D-amino acid (FDAA) TAMRA-D-Lysine (TDL) (Hsu, Meng & VanNieuwenhze, 2016; Kuru *et al.,* 2012; Kuru *et al.,* 2015), which can be incorporated into sites of new peptidoglycan synthesis and would allow for visualization of these sites within the exploring hyphae. We labelled classical vegetative (MYM-grown) and exploring hyphae from wild type and Δ*mreB1* strains after 7 days of exploratory growth on YP agar. In quantifying peptidoglycan labelling patterns, we established three categories to describe the labelling characteristics for individual hyphal compartments (from tip to crosswall): (i) hyphae exhibiting strong side, i.e. dispersed, wall labelling, both alone and in conjunction with polar labelling, (ii) hyphae exhibiting polar labelling without any detectable side wall labelling, and (iii) hyphae exhibiting a random pattern of fluorescent puncta spanning the length of a hyphal compartment (**Fig. 4A**; **Fig. S5A**). We also noted that there were many unlabeled hyphae present in both wild type and *mreB* mutant samples (**Fig. S5B**), suggesting that the TDL label did not indiscriminately associate with exploring hyphal walls; we excluded these unlabeled hyphae from our categorization and quantification. As expected for wild type, we observed strong polar labelling (**Fig. 4A**). We found the majority of labelled hyphae also exhibited side wall labelling. In contrast, side wall labelling was almost completely abolished in our *mreB1* mutant strain, with hyphal labelling being equally divided between the polar and random categories (**Fig. 4B**). This same trend was observed for 5-day exploring samples (**Fig. S5C**). These data suggested that MreB1 may be required for dispersed cell wall synthesis in the lateral walls of exploring *S. venezuelae* cultures.

**Figure 4:**
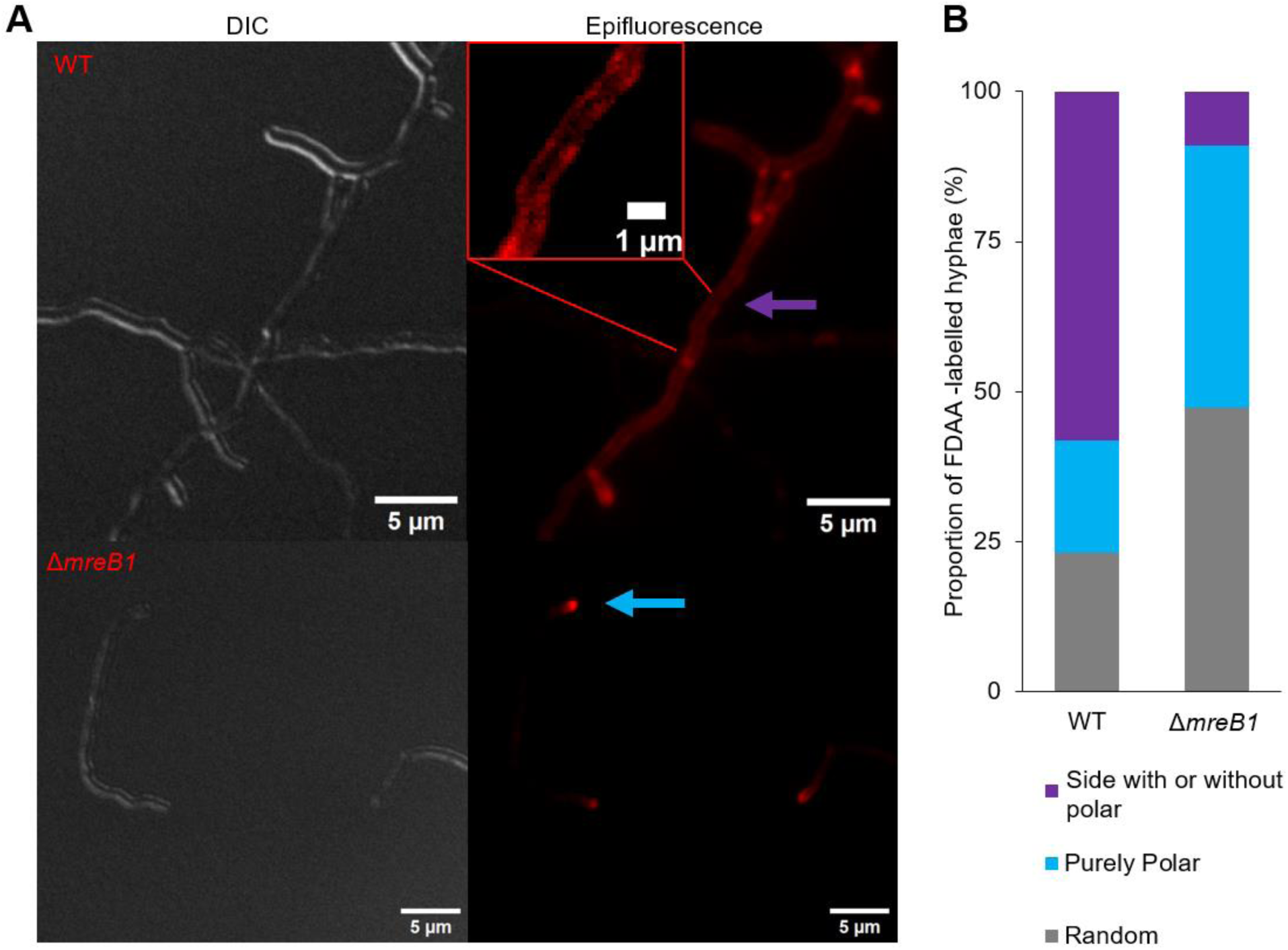
Peptidoglycan insertion patterns of the *mreB1* mutant are distinct from those of wild type. **(A)** Representative DIC and epifluorescent (TRITC/Red) images of wild type (WT) and Δ*mreB1* strains labelled with the fluorescent D-amino acid TAMRA-D-lysine (TDL). Samples were taken from YP-grown cultures after 7 days. Labelled hyphae were sorted into one of three categories: polar and/or side labelled (purple arrow), solely polar labelled (blue arrow) (25 ms exposures). **(B)** Stacked-bar graph indicating the proportion of TDL-labelled exploring hyphae in the groups described in (A). WT: n = 255 labelled hyphae; Δ*mreB1*: n = 336 labelled hyphae. Measured samples were obtained from two independent biological replicates.

### MreB1 localizes to hyphal side walls and exhibits directional movement during exploration

Given the profound decrease in side wall labelling observed for our *mreB1* mutant, we wanted to determine whether MreB1 localized to the lateral walls of exploring hyphae. We replaced *mreB1* with an MreB1-HaloTag sandwich (SW) fusion-expressing variant in its native chromosomal locus and assessed the growth of the resulting strain (**Fig. 5A**; **Video S6**). We found the fusion-expressing strain explored like wild type, suggesting that the fusion was functional (**Fig. 5A**). We next assessed its localization following the addition of the Halo substrate JaneliaFluor 549, compared with a negative control strain expressing the native (untagged) MreB. In classical growth conditions, we observed strong fluorescence intensity associated with sporulation septa and spore walls (**Fig. 5A**). This was consistent with previous MreB localization experiments conducted in *S. coelicolor* (Mazza *et al.,* 2006; Heichlinger *et al.,* 2011) suggesting that the MreB1-HaloTag(SW) protein was localizing as expected.

**Figure 5:**
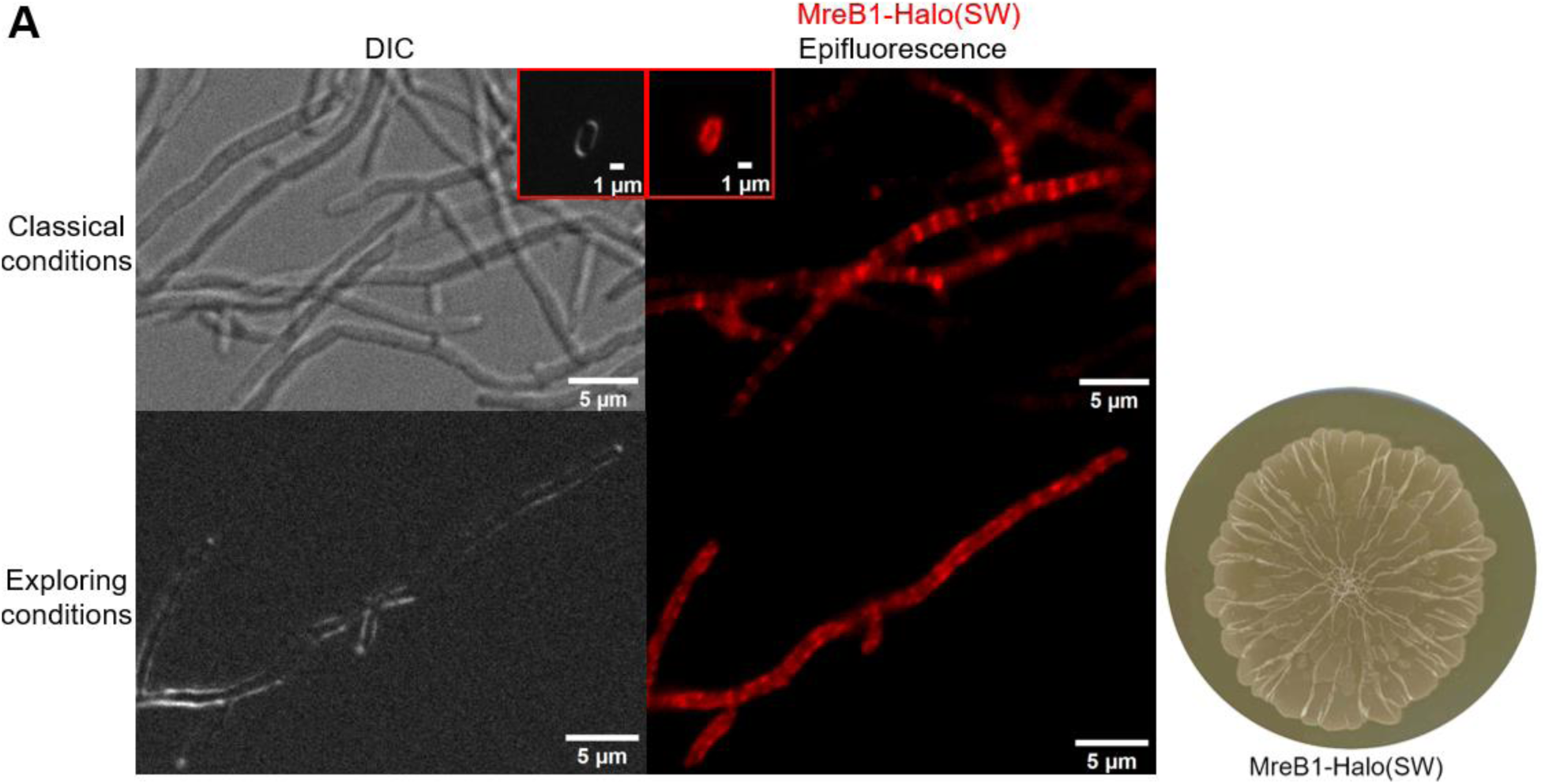
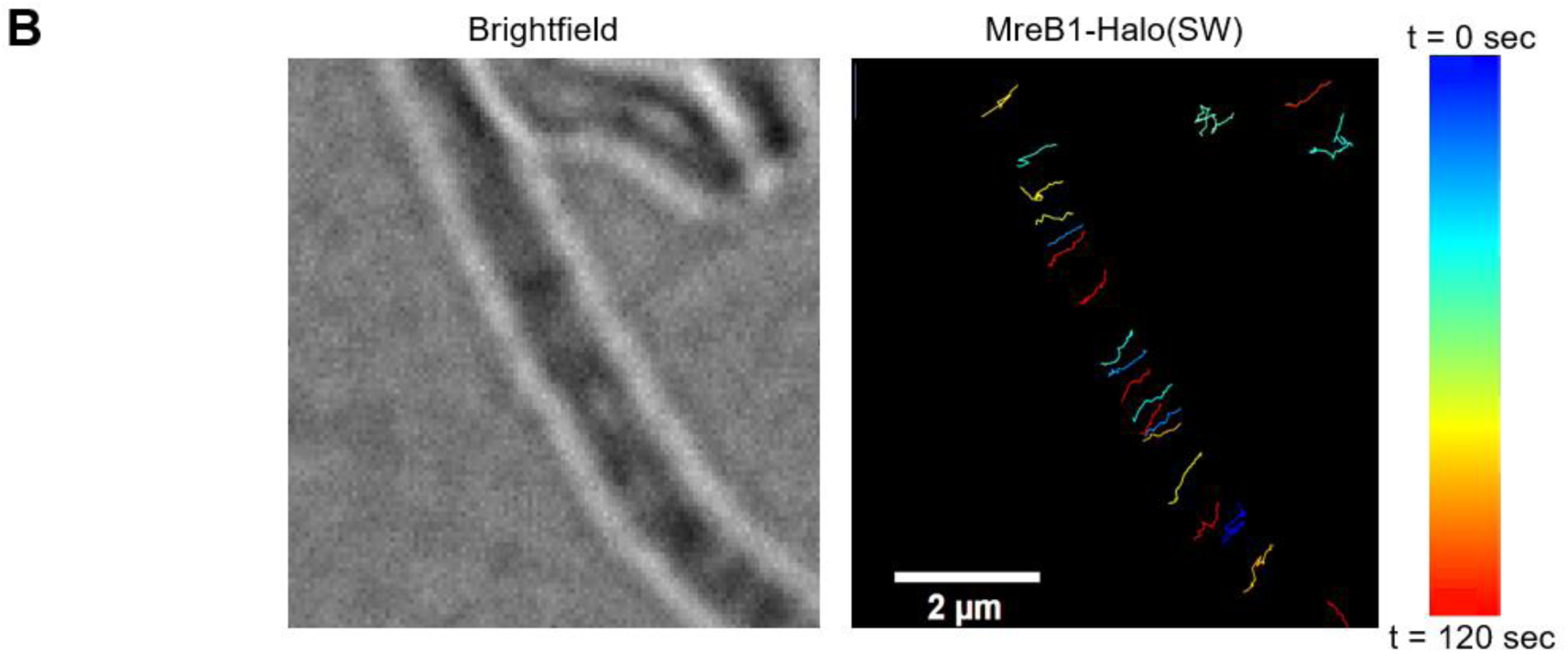
MreB1 exhibits directional movement in *S. venezuelae*. **(A)** Representative DIC and epifluorescent (TRITC/Red) images of *S. venezuelae* expressing a functional MreB1-Halo(SW) fusion (where this fusion has replaced the native copy of *mreB1* on the chromosome). Top left panels: cells grown overnight in liquid MYM (classical conditions; 30 ms exposure) or for 5 days on solid YP plates (exploring conditions; 40 ms exposure). Classical conditions inset: spores expressing MreB1-Halo(SW) (24 ms exposure). Right: exploring MreB1-Halo(SW)-expressing colony grown on solid YP for 10 days. **(B)** Liquid YP-grown MreB1-Halo(SW)-expressing hyphae labelled with Janelia Fluor 549, with MreB being tracked using TrackMate software (Tinevez *et al.,* 2017; Ershov *et al.,* 2022). Tracks were filtered by track length and colour coded based on the time at which tracking for each focus began. Source video (**Video S8**) and unprocessed TrackMate source video (**Video S9**) are available in the supplemental information.

Localizing MreB in an analogous way in exploring cultures proved to be technically challenging, so we sought to identify growth conditions that mimicked exploration conditions but allowed for more effective labelling. We compared the growth of wild type and *mreB1* mutants in liquid YP (exploration-equivalent) and liquid MYM (classical growth-equivalent) medium. We found that in liquid YP, wild type strains grew to much higher OD600 values than during growth in MYM medium, and as noted previously (Jones *et al*., 2017), the pH of the medium rose in a way that was analogous to that seen during exploration. We further observed that during growth in YP medium, the *mreB1* mutant strain exhibited a growth defect relative to wild type, which was not observed in liquid MYM (**Fig. S6A**). Moreover, the Δ*mre* strain also displayed a strong growth defect in liquid YP, and the defective growth trajectories of both the Δ*mreB1* and Δ*mre* strains were improved with supplementation of 25 mM MgSO4, while the wild type remained unaffected (**Fig. S6C**). Finally, we tested whether these liquid YP-grown strains exhibited side wall labelling using the fluorescent TDL substrate, and found that it mirrored what we had seen for YP agar-grown exploring cultures (**Fig. S5D**; **Fig. 4**).

We therefore proceeded to use liquid YP-grown cultures to assess MreB localization. We observed a localization pattern that extended the length of the exploring hyphae, reminiscent of the side wall labelling pattern observed when labelling with the fluorescent D-amino acids during exploration (**Fig. 4 and 5A**). The wild type *S. venezuelae* strain not expressing a HaloTag and labelled with JaneliaFluor 549 was used as a negative control, and showed no hyphal-specific labelling (**Fig. S6B**). To further test the involvement of MreB1 in dispersed wall synthesis in exploring *S. venezuelae*, we acquired timelapse images of MreB1-Halo(SW) during growth in liquid YP. We observed directional movement MreB1, typically perpendicular to the long axis of the hyphal filaments (**Fig. 5B**; **Video S7; Video S8; Video S9**). Importantly, this movement could also be observed for YP agar-grown exploring cells (**Video S10**). These observations supported a role for MreB1 in directing dispersed peptidoglycan synthesis in the lateral walls of rapidly growing cultures.

### MreB is not confined to the sporulating actinobacteria and is conserved throughout the phylum

Historically, MreB has been reported to be either absent from the actinobacteria, or confined to the spore-forming genera where its function is tied to spore wall maturation (Mazza *et al.,* 2006; Heichlinger *et al.,* 2011; Kleinschnitz *et al.,* 2011). Our work here suggests that MreB can also make important contributions to dispersed wall synthesis during the active growth of *Streptomyces* hyphal filaments. Given this functioning of MreB beyond sporulation, we wondered whether *mreB* was indeed confined to the spore-forming actinobacteria, or whether it may be a more ancestral gene within the this phylum. To investigate this, we searched for MreB-encoding actinobacteria, and found many representatives that were not predicted to form spores, although the presence of *mreB* was only broadly conserved within spore-forming genera. We generated a phylogenetic tree of 15 diverse MreB-encoding actinobacteria members, including 5 spore-formers and 10 non-spore formers (**Fig. 6**), and examined the genetic locus surrounding *mreB* in each. In all but two instances, MreB was encoded alongside fellow elongasome proteins MreC and MreD, an organization that is also seen for the model organisms *E. coli* and *B. subtilis*. Many of these regions further mirrored what was found in *S. venezuelae*, including genes encoding a putative class B penicillin-binding protein (PBP) and transglycosylase (RodA) (**Fig. 6**). This suggested that either *mreB* and its associated operon had been acquired multiple times by genera throughout the actinobacteria, or more likely, that the *mreB* operon was present in the founding members of this phylum, and that the *mreB* operon and/or select genes within it had been lost over time.

**Figure 6:**
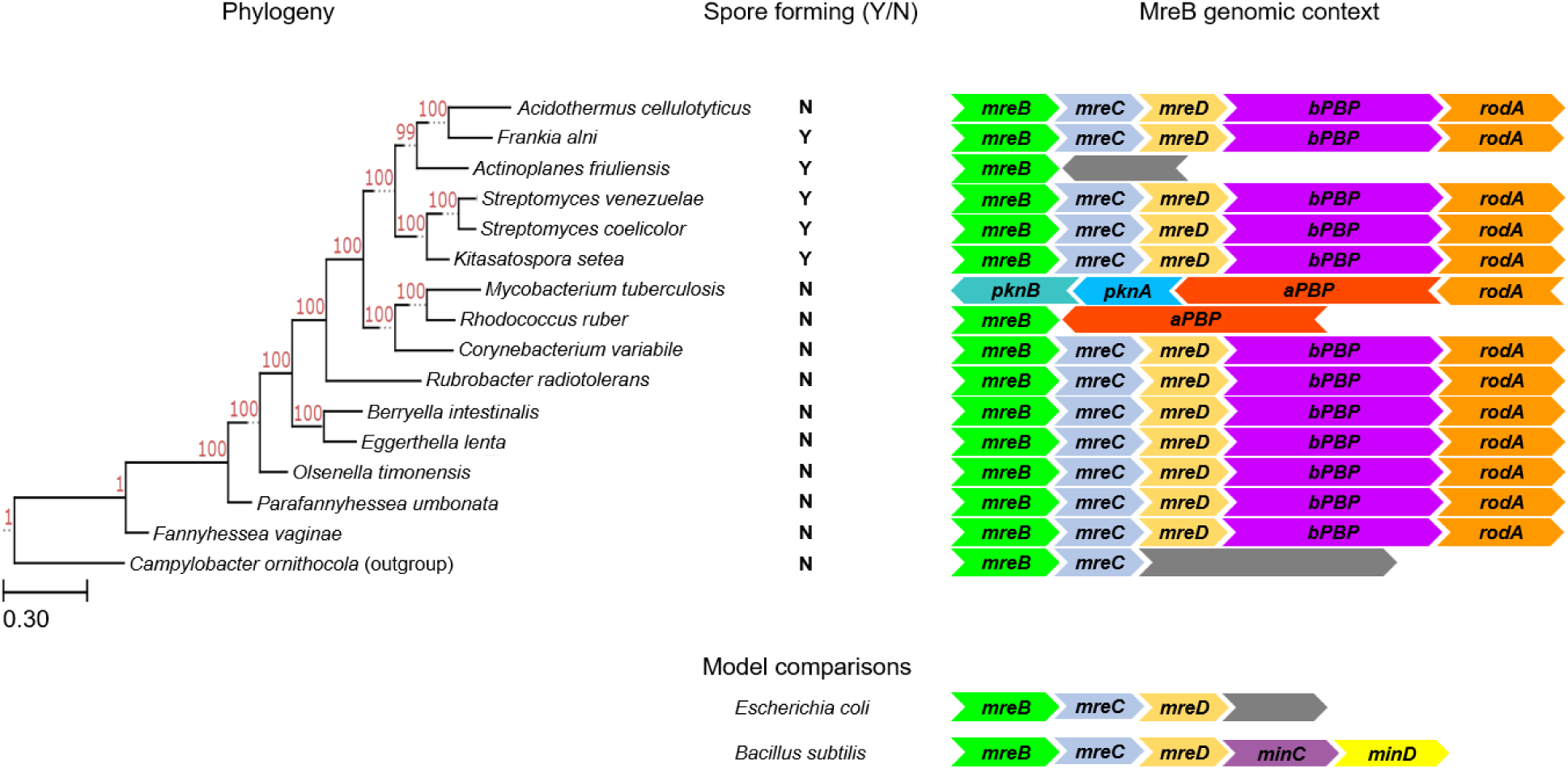
MreB is encoded throughout the Actinobacteria. **(Left)** Phylogenetic tree of MreB-encoding Actinobacteria, using *Campylobacter ornithocola* as an outgroup. Support values (red) of each branch are based on 1000 resampled trials. The scale bar (labelled 0.3) represents substitutions per nucleotide position. AutoMLST (Alanjary, Steinke & Ziemert, 2019) was used for tree construction. **(Middle)** Indication of whether the associated species on the left is known/expected to form spores (Y) or is not considered to be a spore former (N). **(Right)** Schematic (not to scale) of the genomic context of *mreB* genes in the species presented within the tree to the left; the genetic organization around *mreB* in the model organisms *E. coli* and *B. subtilis* is presented below. Proteins commonly encoded adjacent to MreB include MreC, MreD, a class B PBP (bPBP), the RodA transglycosylase, and less commonly, a class A PBP (aPBP), and the Min proteins (MinC and MinD) in *B. subtilis. Mycobacterium tuberculosis* does not encode MreB, therefore the genomic context for RodA is displayed.

## Discussion

In bacteria, growth can occur by incorporating cell wall material at the cell poles, along the lateral walls, or at the division plane. While the vast majority of bacteria undergo cell division (and thus cell wall synthesis at the division plane), until now, polar growth and dispersed wall growth have been considered mutually exclusive systems, with bacteria employing one or the other but not both. Here, we show that in *S. venezuelae*, exploratory growth requires a combination of DivIVA-driven polar growth (polarisome) and MreB-mediated dispersed side wall synthesis (elongasome), and this has implications for the cell wall architecture of this bacterium over the course of an exploratory growth cycle. Conventional polarisome-mediated growth in exploring *S. venezuelae* was apparent from the robust labelling of exploring hyphal cell poles with FDAAs, which marked sites of new peptidoglycan synthesis. Under rapid growth conditions like those associated with exploration, the importance of MreB-mediated elongasome activity was supported by multiple lines of evidence, including (i) the incorporation of FDAAs into hyphal side walls, (ii) the localization of MreB to the lateral walls, (iii) the directional movement of MreB around the circumference of hyphal cells, (iv) the exploration defects and cell lysis associated with the loss of MreB, and (v) the altered peptidoglycan composition associated with an *mreB* mutant specifically under growth conditions where its loss has phenotypic consequences.

The MreB-dependent change in cell wall width that accompanied progression through an exploration cycle was also notable and unexpected. Elegant work in *E. coli* has shown cell width can change in response to environmental conditions, but that the width of the wall remains constant (Tropini *et al.,* 2014). Our results here suggest that in the early stages of exploration, the cell wall is thinner, perhaps indicating that the onset of exploration involves a trade-off between enhanced DivIVA-mediated cell extension and reduced peptidoglycan thickness. It appears, however, that this trade-off is not viable in the long term, as the inability to reinforce the cell wall in the absence of MreB was correlated with increased cell lysis. In other systems where MreB drives cell wall synthesis, *mreB* depletion leads to cell lysis after several generations of defective peptidoglycan synthesis (*e.g., B. subtilis* and *Caulobacter crescentus*) (Gitai *et al.,* 2005), consistent with our observations here. The apparent MreB-mediated side wall reinforcement seen here is further reminiscent of the growth of plant cells, which are initially synthesized with a thin, flexible wall that allows for cell expansion, followed by deposition of a secondary cell wall to provide strength and rigidity to the cells (McKenna *et al*., 2009; Derbyshire *et al*., 2007).

*Streptomyces* have long been considered outliers within the better-studied actinobacteria (*e.g. Mycobacterium, Corynebacterium*) in encoding both DivIVA and MreB, and it had been assumed that MreB functioned solely during spore formation. However, we found that both MreB and its associated elongasome members were broadly distributed throughout the actinobacteria. This suggests that the ability to couple polar growth with dispersed wall synthesis may be widespread within this group. While dispersed wall synthesis has not been reported for other actinobacteria, recent work has revealed a role for the RodA protein in repairing lateral wall damage in *Mycobacterium smegmatis* (García-Heredia *et al.,* 2018). In this system, the sensing of cell wall damage leads to side wall localization of RodA, and an associated redeployment of the cell wall biosynthetic machinery from the poles to the sides to enable cell wall repair. In *Streptomyces*, the cell wall damage response is mediated by the alternative sigma factor σ^E^. Notably, σ^E^ function is essential for exploration (Shepherdson et al., 2024), and previous work in *S. coelicolor* has shown that *mreB* is under σ^E^ control (Paget *et al.,* 1999; Paget, Leibovitz & Buttner, 1999; Hong, Paget & Buttner, 2002; Tran *et al.,* 2019). These observations collectively suggest there may be evolutionary connections linking the lateral wall repair capabilities of the mycobacteria, and the dispersed wall synthesis capabilities of *S. venezuelae*. The biosynthetic components of the *Streptomyces* polarisome have not been defined, and thus it remains to be determined whether there are shared elements between the polarisome and the elongasome (as appears to be the case in the incomplete mycobacterial system), or whether these are entirely discrete cell wall biosynthetic units. Similarly, the signal that promotes MreB recruitment and activation in *S. venezuelae* has yet to be identified; however, it is tempting to speculate that it may relate to the cell wall damage response.

Our work here reveals that the cell wall biosynthetic capabilities of bacteria can be far more flexible than previously appreciated, with *Streptomyces* bacteria having the ability to simultaneously employ polarisome-and elongasome-mediated cell wall synthesis. Furthermore, the presence of equivalent systems throughout the actinobacteria suggests that this ability may be more widely distributed.

## Materials and Methods

### Strains, plasmids, media, and culture conditions

Strains, plasmids, and primers used in this study are listed in Table S3. *S. venezuelae* NRRL-B-65442 (Gomez-Escribano *et al*. 2021) was grown on solid (2% agar) MYM (1.0% malt extract, 0.4% yeast extract, 0.4% maltose) for spore stock generation, and grown overnight for vegetative microscopic analysis and isolation of peptidoglycan. Exploration experiments were conducted using solid (2% agar) YP (1% yeast extract, 2% peptone) or YPD (1% yeast extract, 2% peptone, 2% D-glucose) with the yeast *S. cerevisiae* (YPDy). For the magnesium supplementation experiment, YP medium was supplemented with 25 mM MgSO4. All growth incubations were carried out at 30°C, except for overnight growth on solid MYM (between 12-16 h), which was conducted at room temperature to slow growth and development. For strain growth comparisons on YP/YPDy, 5 μL of titred spore stocks (normalized to lowest CFU/mL for spore titres being compared) were spotted directly onto YP/YPD agar plates. For YPDy exploration experiments, 5 μL of an overnight *S. cerevisiae* (yeast) culture (grown at 30°C in liquid YPD) were spotted immediately adjacent to 5 μL of the *S. venezuelae* spore stock. For all agar-grown cultures, 40 mL of YP/YPD were poured (to account for the incubation periods of up to 20 days required for exploration experiments), while all MYM plates were 25 mL (where growth periods did not extend beyond one week). All exploration plates were incubated for up to 12 days (YP) or 20 days (YPDy). Videos were gathered of strains growing at 30°C, using the Epson Perfection V800 Photo scanner (Epson, Nagano, JA) which had been programmed to acquire one image per hour. Images were then compiled into time-lapse video formats. The OD_600_ growth curves (**Fig. S6A**) performed for wild type and *mreB1* mutant strains involved normalizing overnight cultures to a starting OD_600_ of 0.192 in 50 mL of either liquid MYM or YP media in 250 mL baffled Erlenmeyer flasks. The cultures were then grown at 30°C and 200 rpm shaking, for up to 18 hours. For the OD_600_ growth curves in **Fig. S6C**, cultures were grown in 1 mL of liquid YP medium with or without supplementation of 25 mM MgSO_4_ in a 48-well plate, with continuous shaking at 30°C. OD_600_ values were measured every 15 min, and each data point represents the average of three individual biological replicates using the Cytation3 plate reader (BioTek, VT, USA). Each sample was normalized to a starting OD_600_ value of 0.05 from an MYM-grown liquid overnight culture.

*E. coli* strains were grown with appropriate antibiotic selection in or on LB (lysogeny broth), SOB (super optimal broth), or Difco nutrient agar plates. DH5α and ET12567/pUZ8002 strains were grown at 37°C, while BW25113/pIJ790 was grown at 30°C or 37°C.

### Construction of *S. venezuelae* mutant, complementation, and translational fusion strains

Gene deletions were generated using the ReDirect method (Gust *et al.,* 2003). The coding sequences for *vnz_11695* (*mreB1*), *vnz_11715* (*mreB2*), and *vnz_11675-vnz_11695* (*mre* operon) were replaced in the Sv-6-F06 cosmid vector with an *aac(3)IV-oriT* apramycin resistance cassette. For the Δ*mreB1/*Δ*mreB2* double mutant strain, the coding sequence for *mreB1* was replaced with a viomycin resistance cassette, while the coding sequence for *mreB2* was replaced with a marker-less, in-frame deletion, using a cosmid in which the apramycin resistance cassette that had replaced the *mreB2* gene (described above) and was flanked by FLP recognition target (FRT) sites, was removed using FLP-mediated recombination. Modified cosmids were transformed into *E. coli* ET12567 carrying the pUZ8002 helper plasmid that enables conjugation into *S. venezuelae*. Double crossover exconjugants were selected for using apramycin or viomycin selection and deletions were confirmed using diagnostic primer combinations annealing to regions upstream, downstream and internal to the coding sequences. For conjugation of the clean *mreB2* deletion, single crossovers were selected for with kanamycin (where resistance was conferred by a gene in the cosmid backbone) followed by screening for double crossovers (loss of kanamycin resistance) and PCR confirmation.

For complementation, the coding sequences and native promoters were PCR amplified and cloned into the hygromycin resistant gene-containing integrative pMS82 plasmid, which had been further modified to carry a kanamycin resistance gene. Construct integrity was confirmed by sequencing, before being introduced into conjugation-competent *E. coli*. Following conjugation from *E. coli* ET12567 into *S. venezuelae* and selection using nalidixic acid (to select against the *E. coli* donor strain) and kanamycin/hygromycin (to select for gain of the integrative plasmid), successful transfer and integration of the plasmid into the chromosome were confirmed using primers designed to amplify the kanamycin resistance gene.

The MreB1-HaloTag(SW) fusion was constructed using NEBuilder HiFi DNA Assembly (New England Biolabs, MA, USA) using native promoters and cloning into the TOPO2.1 vector conferring kanamycin resistance. Single crossovers were selected for with kanamycin, followed by screening for double crossovers (loss of kanamycin resistance). Successful recombination of the *mreB1-HaloTag(SW)* fusion was confirmed with diagnostic PCR reactions.

Overexpression of *mNeonGreen* for testing cell viability and cell lysis involved cloning the *mNeonGreen* coding sequence behind the constitutive *ermE\** promoter in the kanamycin/hygromycin resistant pMS82 plasmid. The resulting construct was introduced into wild type and mutant strains via the conjugation process described above. All primers used for generating ReDirect deletions, complementation constructs, translational fusions, and overexpression constructs, as well as diagnostic PCR primers used for confirmation of construct presence/mutant creation, are listed in Table S3.

### Transmission and scanning electron microscopy

For both TEM and SEM, samples were fixed with 2% glutaraldehyde (2% v/v) in 0.1 M phosphate buffer (pH 7.4). The samples were rinsed twice in buffer solution, before being post-fixed in 1% osmium tetroxide in 0.1 M phosphate buffer for 1 hour. The samples were then dehydrated through a graded ethanol series (50%, 70%, 70%, 95%, 95%, 100%, 100%).

The final dehydration for the TEM samples was done in 100% propylene oxide (PO). Infiltration with Spurr’s resin was through a graded series (2:1 PO:Spurr’s, 1:1 PO:Spurr’s, 1:2 PO:Spurr’s, 100% Spurr’s, 100% Spurr’s, 100% Spurr’s) with rotation of the samples in between solution changes. The samples were then transferred to embedding moulds filled with fresh 100% Spurr’s resin and polymerized overnight in a 60°C oven. Thin sections were cut using a Leica UCT Ultramicrotome and picked up onto Cu grids. The sections were post-stained with uranyl acetate and lead citrate and viewed using a JEOL JEM 1200 EX TEMSCAN transmission electron microscope (JEOL, Peabody, MA, USA) operating at an accelerating voltage of 80 kV.

For SEM samples, after dehydration in 100% EtOH, samples were critical point dried, mounted onto SEM stubs, sputter-coated with gold and viewed in a Tescan Vega II LSU scanning electron microscope (Tescan, PA, USA) operating at 20 kV.

### Peptidoglycan purification and UPLC-MS analysis

Peptidoglycan samples were analysed as described previously, following the 24-hour protocol with some modifications (Alvarez *et al*., 2016; Kühner *et al*., 2014). In brief, samples were boiled in 5% SDS for 4 hours, after which the resulting sacculi were washed repeatedly with MilliQ water followed by ultracentrifugation (110,000 rpm, 10 min, 20°C). The washed sacculi were then sonicated, and treated with DNase, RNase, and trypsin protease, after which these enzymes were inactivated by boiling. Wall teichoic acids were removed using 1 M HCl. The samples were finally treated with muramidase (100 μg/mL) for 15 hours at 37°C. Muramidase digestion was stopped by boiling, and coagulated proteins were removed by centrifugation for 10 min at 14,000 rpm. The resulting wall suspensions were then pH adjusted to 8.5-9.0 with sodium borate buffer, after which sodium borohydride was added to a final concentration of 10 mg/mL. After reduction for 30 min at room temperature, the sample pH was adjusted to pH 3.5 with orthophosphoric acid.

UPLC analyses of muropeptides were performed using a Waters UPLC system (Waters Corporation, MA, USA) equipped with an ACQUITY UPLC BEH C18 Column, 130 Å, 1.7 μm, 2.1 mm × 150 mm (Waters Corporation, MA, USA) and a dual wavelength absorbance detector. Muropeptides were separated at 45°C using a linear gradient from buffer A (formic acid 0.1% in water) to buffer B (formic acid 0.1% in acetonitrile) in a 25 min run, with a 0.50 mL/min flow. Muropeptide elution was detected at 204 nm. Relative total peptidoglycan amounts were calculated by comparing the total intensities of the chromatograms (total area) from three biological replicas normalized to the same initial biomass and extracted with the same volumes. Muropeptide quantification was based on their relative abundances (relative area of the corresponding peak).

### FDAA labelling

*S. venezuelae* cells (scraped from solid MYM or YP plates) were incubated at room temperature in a 400 μL solution of 0.4 μM TAMRA-D-Lysine (TDL) (Kuru *et al.,* 2015) for 5 min. Cells were then fixed in 70% ethanol for 15 min at -20°C, after which cells were washed four times with phosphate buffered saline (pH 7.4). Samples obtained from liquid culture were diluted 1/10 prior to FDAA labelling. All other steps were carried out as described above.

### Fluorescence microscopy and colony visualization

Fluorescence microscopy was performed using 1% agarose pads and with an A1R HD25 inverted confocal microscope (NIKON, Tokyo, JA) TRITC [Excitation filter 540/25; TDL and MreB1-Halo(SW)] with ×60 magnification using an oil immersion objective. All exposure times were calibrated using negative control samples (unlabelled wild type *S. venezuelae* for TDL labelling, and wild type *S. venezuelae* labelled with Janelia Fluor 549 for MreB1-Halo(SW) labelling) to eliminate background and autofluorescence. Experiments comparing mNeonGreen fluorescence intensity images were taken using the Typhoon FLA-9500 laser scanner (GE Healthcare, IL, USA) in the AlexaFluor-488 channel (wavelength range: ≥510 nm), with four edge-associated, non-wrinkle-related regions being quantified per exploring colony. Still images of live cells encoding MreB1-HaloTag(SW) were captured following labelling with 1 μM Janelia Fluor 549 (Promega, WI, USA) for 1 hour in phosphate-buffered saline, after which they were washed three times (30 min per wash) in phosphate-buffered saline.

### Timelapse TIRFM

Timelapse TIRFM images were taken with 300 ms exposure using the 561 nm laser line at 2 s intervals. All components were prewarmed to 30°C including MatTek dishes, agarose pads and media. Samples were labelled for 30 min at 30°C in liquid YP with a 250 nM concentration of Janelia Fluor 549 dye.

### Image analyses

All images were analyzed using Fiji software (Schindelin *et al.,* 2012). The analyses included cell wall thickness measurements of TEM micrographs, quantification of FDAA TDL labelling, quantification of relative mNeonGreen fluorescence intensity, and exploring colony surface area measurements. Brightness/contrast of whole colony mNeonGreen images were adjusted to the minimal level needed to eliminate background fluorescence from a wild type empty-vector *S. venezuelae* control. TrackMate (Tinevez *et al.,* 2017; Ershov *et al.,* 2022) software was used to follow the directional movement of MreB1-Halo(SW). Tracks presented were filtered for overall track length, and the unedited video is available in the supplemental information.

### Phylogenetic analyses

The AutoMLST program (Alanjary, Steinke & Ziemert, 2019) was used to generate the phylogenetic tree. NCBI accession numbers used for analysis were as follows; *Acidothermus cellulolyticus* (NC_008578.1), *Actinoplanes friuliensis* (NC_022657.1), *Berryella intestinalis* (NZ_CP009302.1), *Corynebacterium variabile* (NC_015859.1), *Eggerthella lenta* (NZ_CP089331.1), *Fannyhessea vaginae* (NZ_CP065631.1), *Frankia alni* (NC_008278.1), *Kitasatospora setea* (NC_016109.1), *Mycobacterium tuberculosis* (NC_000962.3), *Olsenella timonensis* (NZ_LT635455.1), *Parafannyhessea umbonata* (NZ_LT629759.1), *Rhodococcus ruber* (NZ_CP044211.1), *Rubrobacter radiotolerans* (NZ_CP007514.1), *Streptomyces coelicolor* (NZ_CP042324.1), and *Streptomyces venezuelae* (NZ_CP018074.1).

## Acknowledgements

We would like to thank members of the Elliot lab, Yves Brun (Université de Montréal), Joseph McCormick (Duquesne University), and Susan Schlimpert (John Innes Centre) for helpful discussions, Michelle Williams (McMaster University) and Daniel Henthorn (Harvard University) for microscopy assistance, and Gerard Wright and Kalinka Koteva (McMaster University) for their generous gift of the TDL FDAA. This work was supported by a Natural Sciences and Engineering Research Council (NSERC) Discovery Grant (to MAE). MPZ and CRB were supported by an Ontario Graduate Scholarship, while SEJ was supported by an NSERC Vanier Scholarship.

---

**Tables S1&2 – Independent Word File**

**Table S3:**
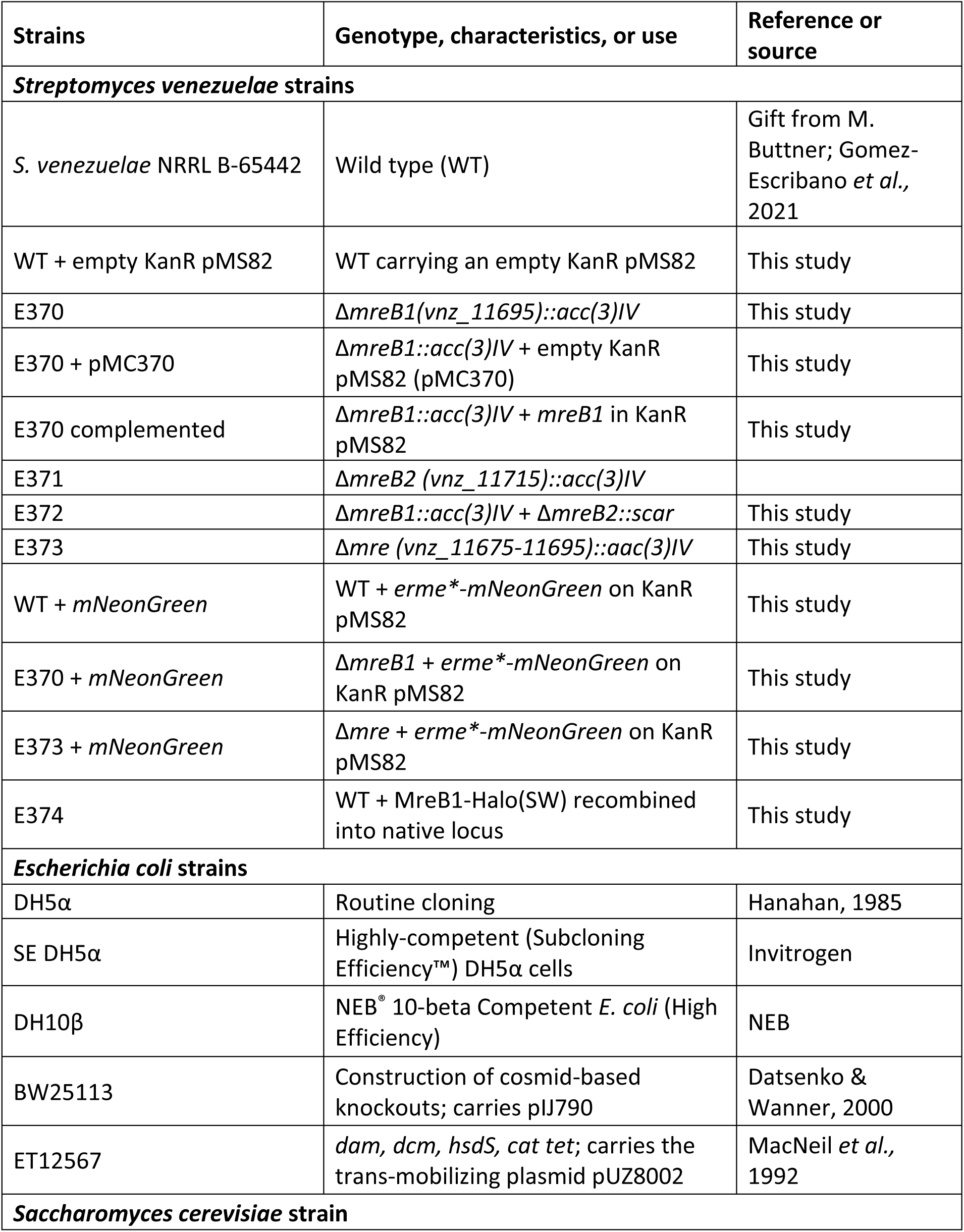
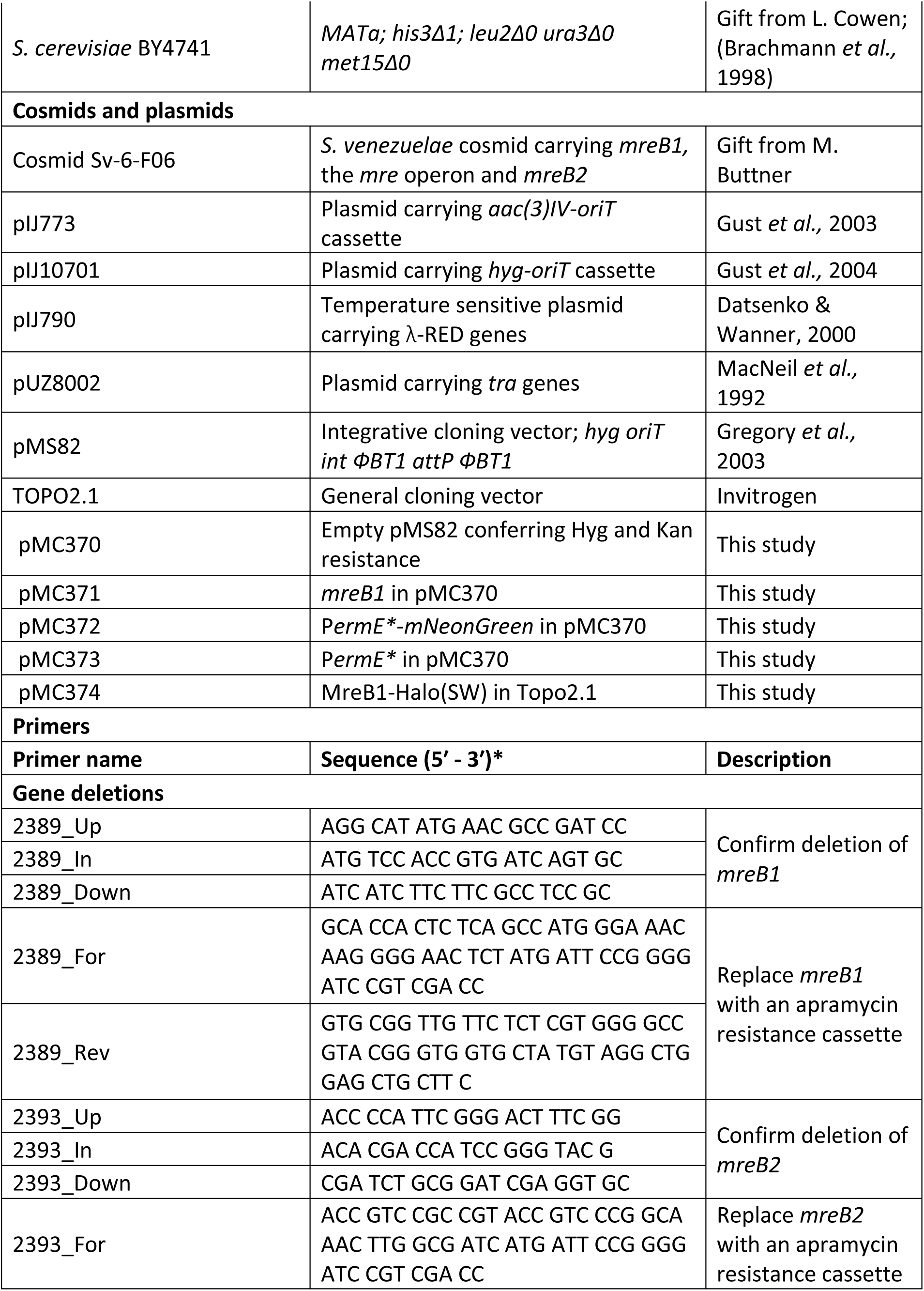
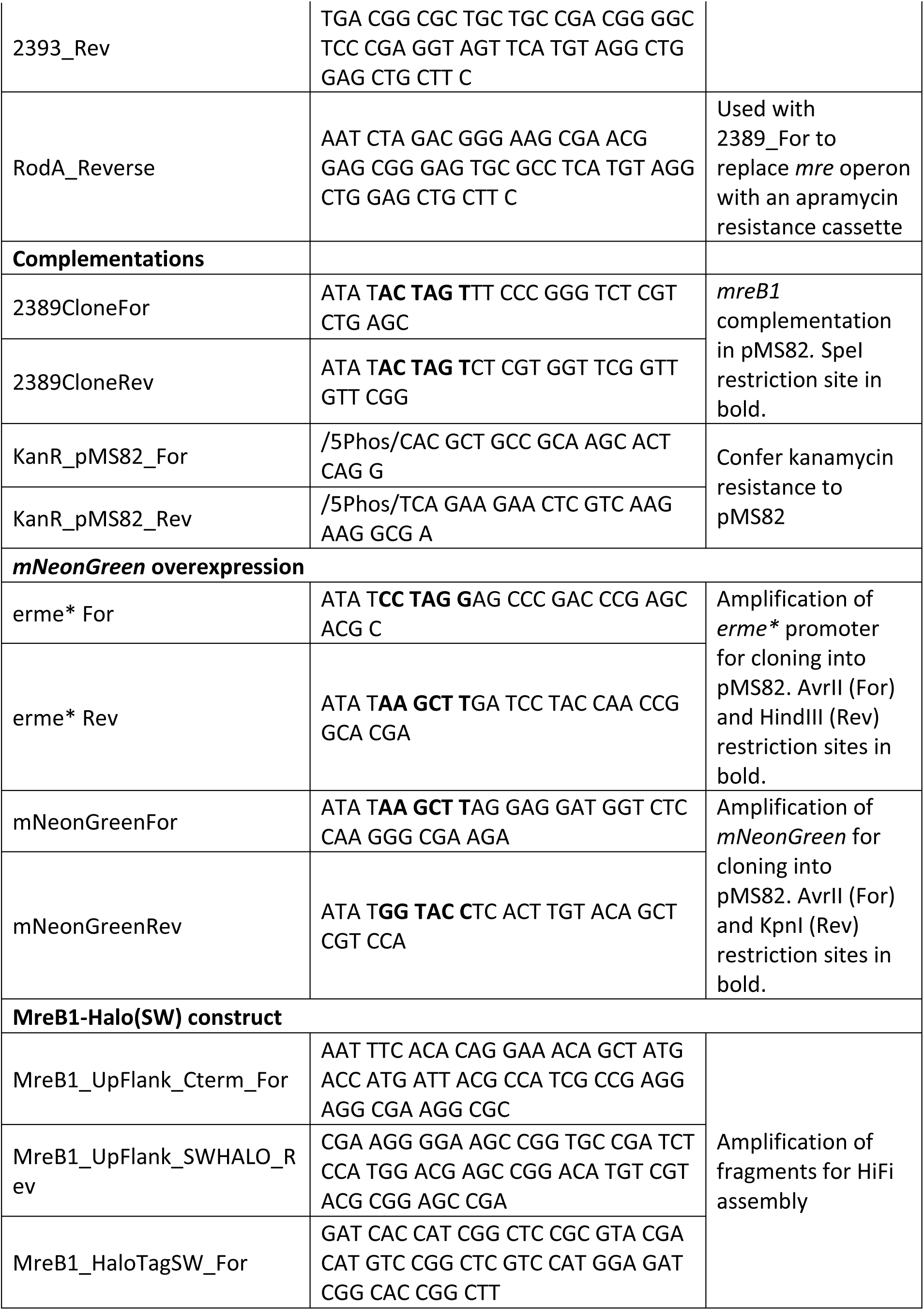
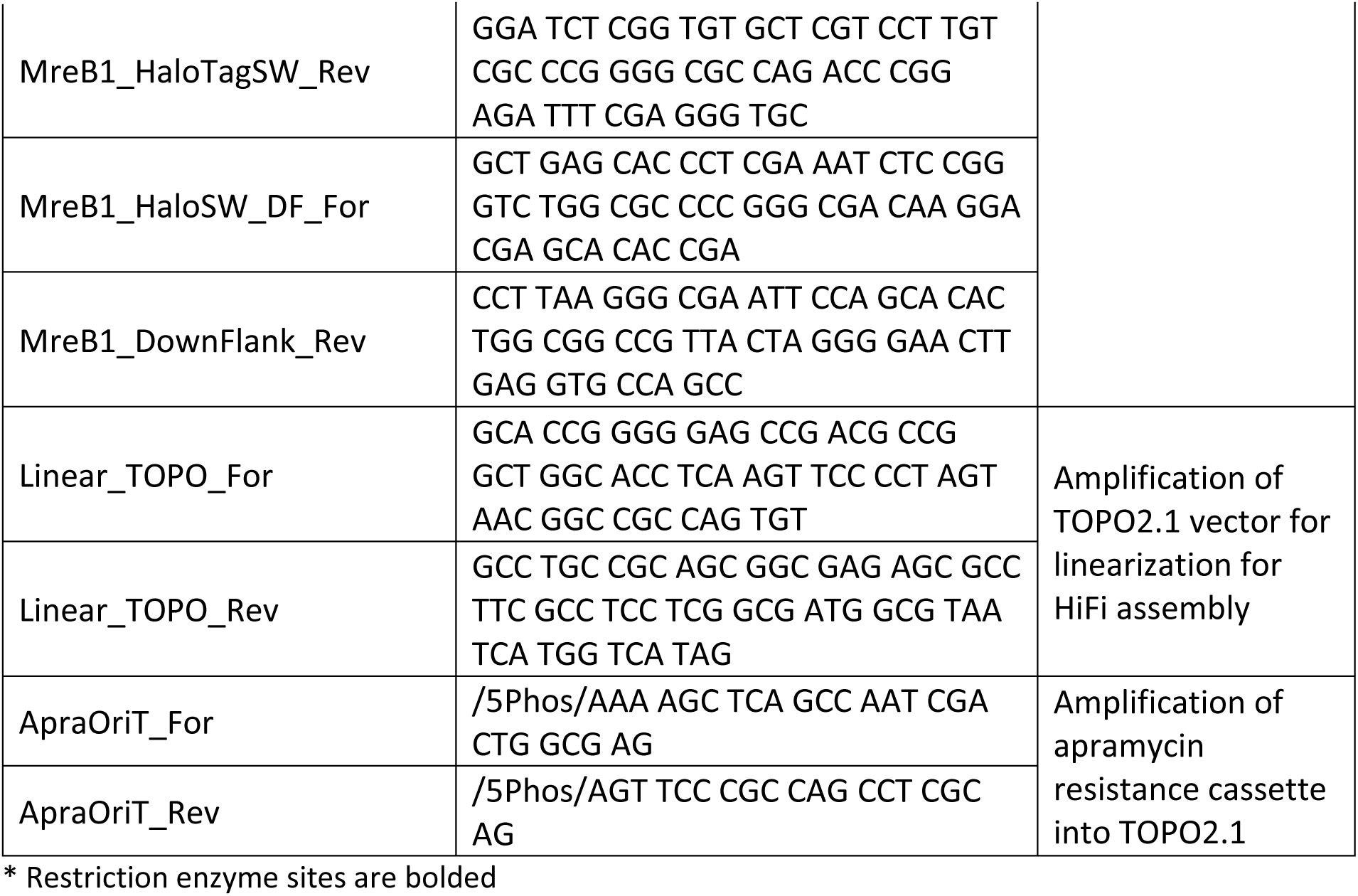
Strains, plasmids and primers used in this study.

## Supplementary data

**Figure S1:**
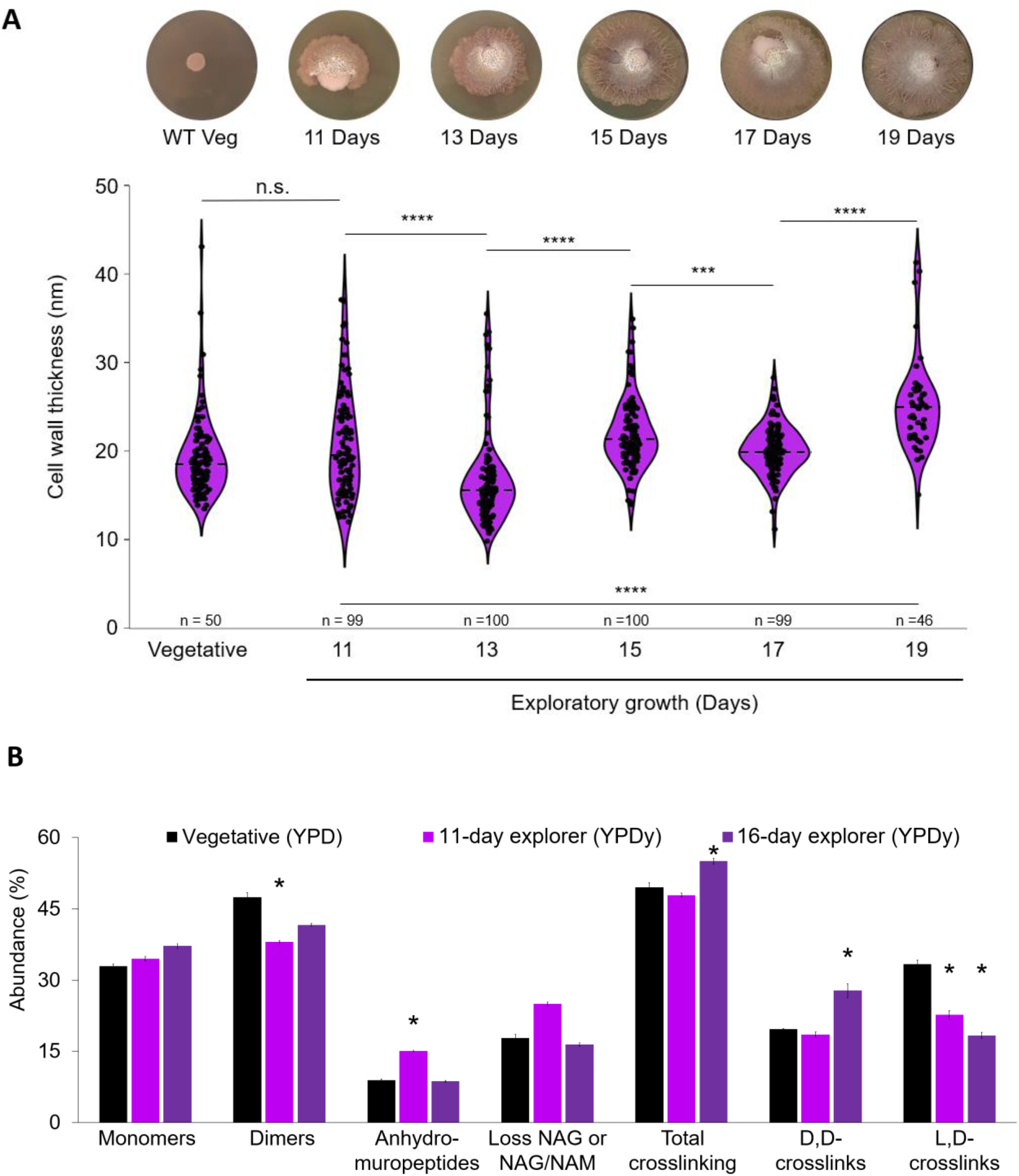
YPDy explorers show similar trends as YP explorers in their cell wall composition compared with vegetatively grown cells. **(A)** Violin plot of hyphal cell wall thickness of vegetative (MYM) and YPDy explorers over time, as determined by measuring transmission electron micrographs using ImageJ. Three measurements taken for each hypha, and averaged. Dashed lines denote median values. Above the plots are representative images of vegetative/exploring colonies from the time points indicated. Statistical significance was determined via Mann-Whitney tests. **(B)** The abundance of total crosslinking, D,D-crosslinks and L,D-crosslinks upon isolation and UPLC-MS analysis of digested peptidoglycan from wild type samples grown on YPD (vegetative) or YPDy exploration medium. Asterisks indicate statistically significant differences between exploring and vegetative conditions (* = P value < 0.05, ** = P value < 0.01, *** = P value < 0.001, **** = P value < 0.0001; as determined by Student’s t-tests). Error bars indicate standard error. Data presented are the average values of two replicates (YPD, 16-day YPDy) of three replicates (11-day YPDy).

**Figure S2:**
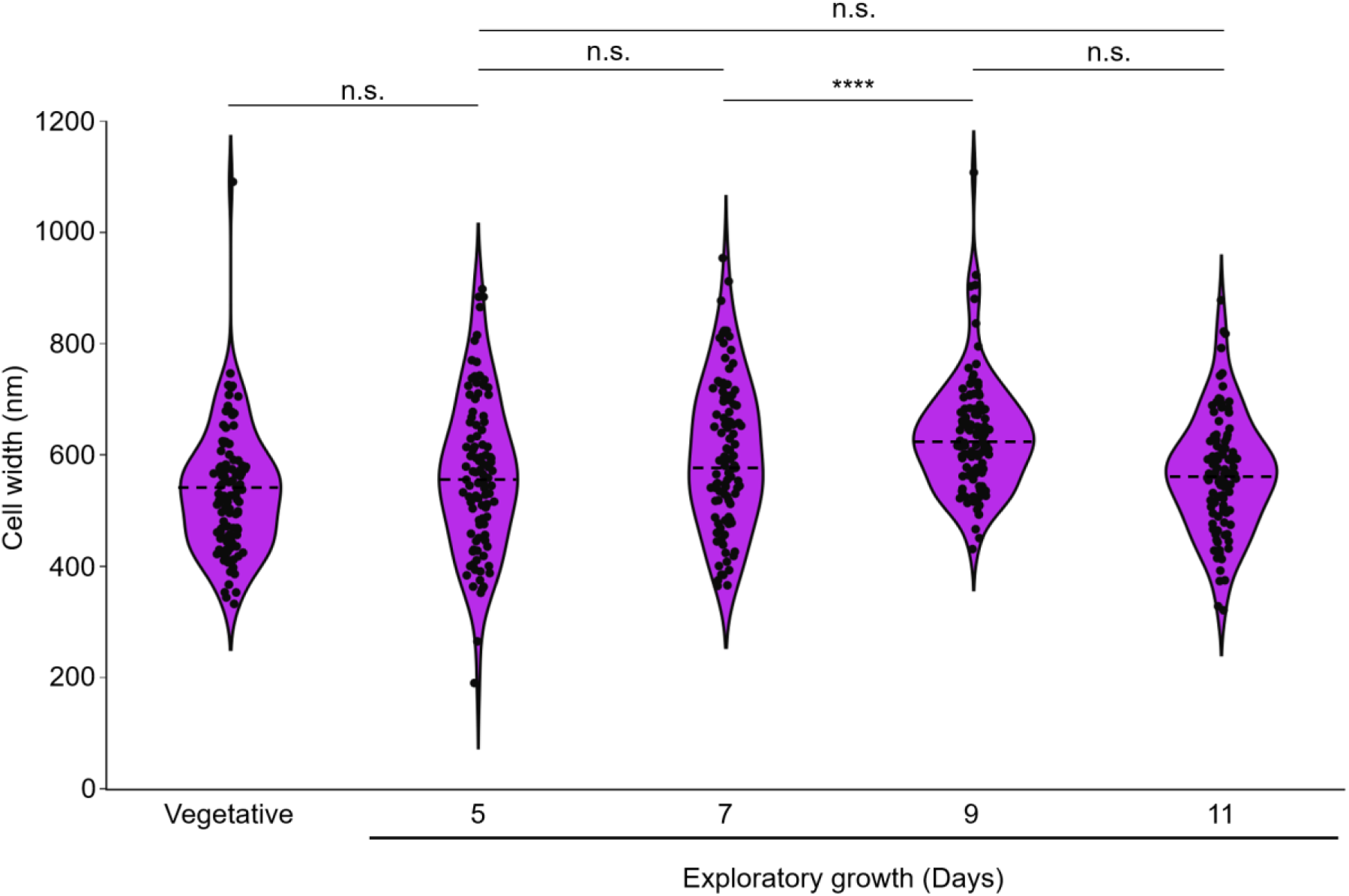
Cell width of YP-grown exploring cells and vegetative (MYM) hyphae do not exhibit the same dramatic changes over time as the cell walls. Samples measured here are the same as those presented in Figure 1A, only measured for cell width. Dashed lines denote median values from hyphae obtained using TEM and measured using ImageJ. Asterisks denote statistical significance, as determined by Mann-Whitney tests.

**Figure S3:**
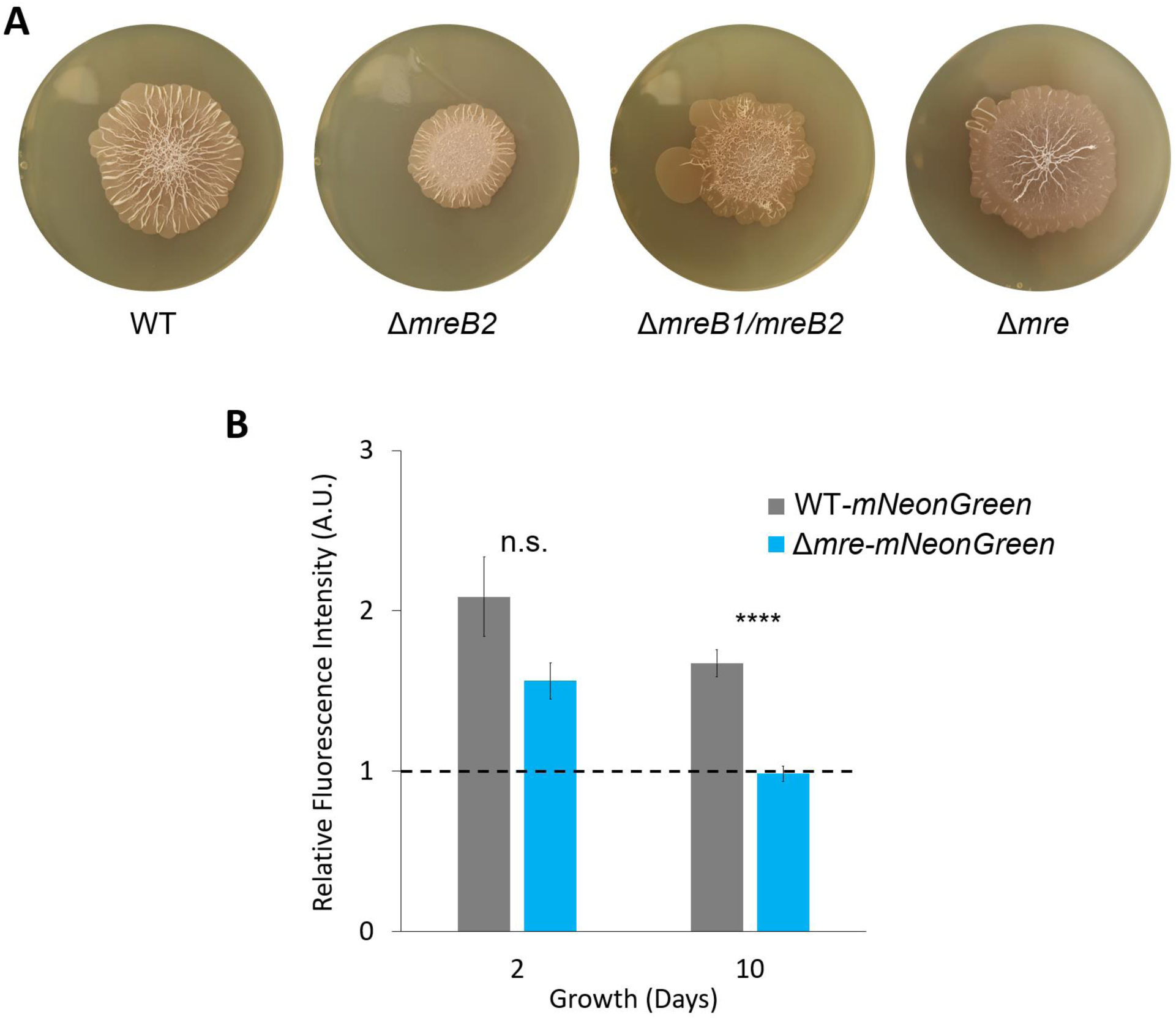
MreB2 does not make a major contribution to exploration but the *mre* operon is important. **(A)** Plate images of wild type, Δ*mreB2,* Δ*mreB1/mreB2* and Δ*mre* strains of *S. venezuelae* grown on YP agar for 10 days. **(B)** Comparison of relative whole-colony fluorescence between wild type and Δ*mre* strains with constitutively expressed *mNeonGreen*. Values presented are relative to a negative control strain lacking *mNeonGreen* and are the average of biological triplicates. Cultures were grown on YP exploration medium. Asterisks indicate statistically significant differences (n.s. = P value > 0.05; **** = P value < 0.0001; as determined by Student’s t-tests).

**Figure S4:**
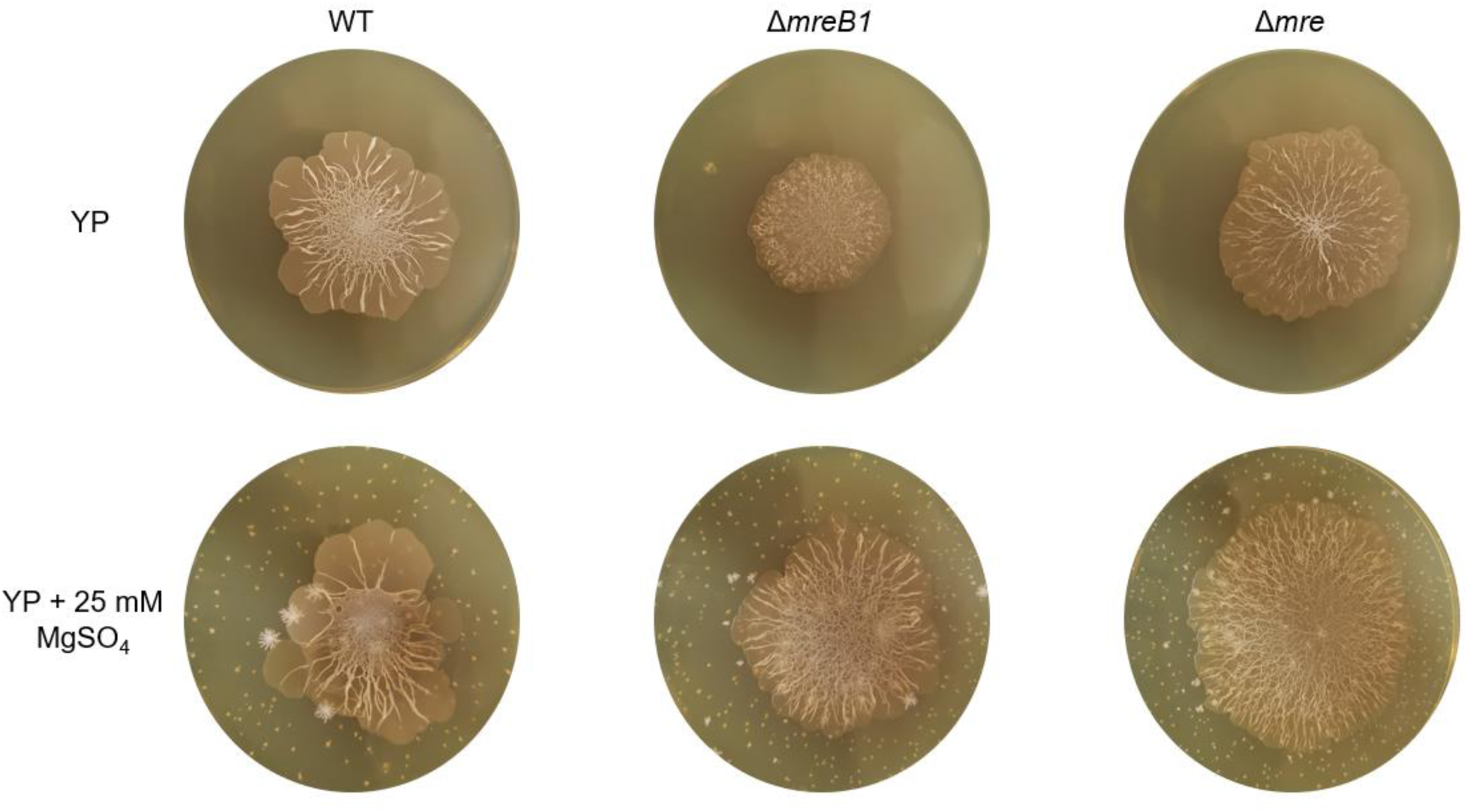
Magnesium supplementation rescues the lysis defect of *mreB1* and *mre* operon mutants. Wild type, Δ*mreB1* and Δ*mre* strains were grown on YP medium (top) and YP medium supplemented with 25 mM MgSO_4_ (bottom) for 10 days. Prolonged growth on YP medium and the corresponding evaporation of growth medium result in the precipitation of MgSO_4_, giving a spotty appearance to the agar plates.

**Figure S5:**
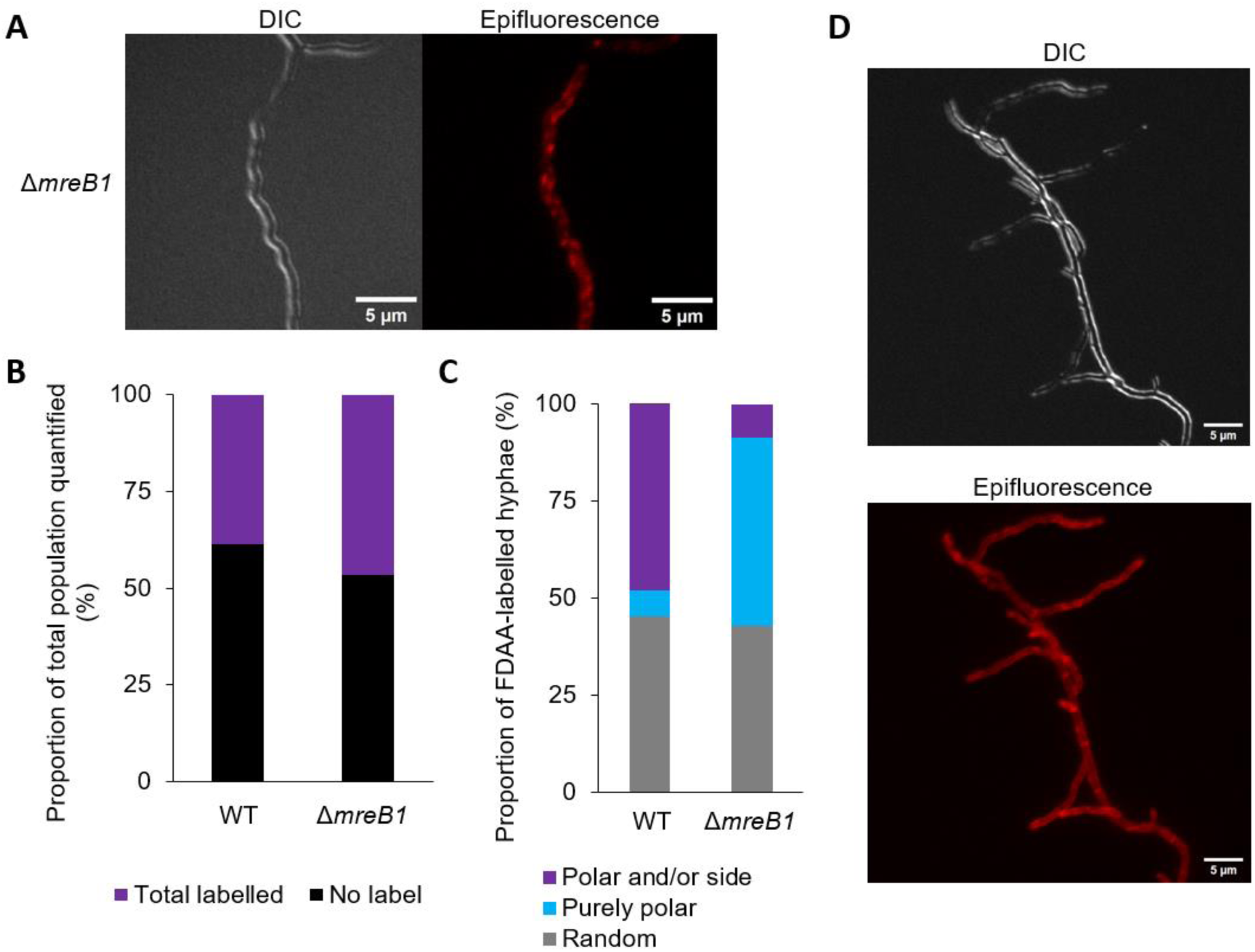
FDAA labelling is specific to a subset of hyphae and reveals that 5-day explorer cells have the same incorporation pattern as 7-day explorer cells. **(A)** Representative DIC and epifluorescence (TRITC/Red) images of the Δ*mreB1* strain labelled with the fluorescent D-amino acid TAMRA-D-lysine (TDL) displaying a random labelling pattern (25 ms exposure) after 7 days of growth on YP medium. **(B)** Stacked-bar graph displaying the proportion of TDL-labelled exploring hyphae (purple) and the total proportion of presumably non-growing/dead hyphae that did not label with TDL (black). Wild type: n = 661 labelled hyphae; Δ*mreB1*: n = 720 labelled hyphae. Samples measured were obtained from two independent biological replicates. **(C)** Stacked-bar graph displaying the proportion of TDL-labelled exploring hyphae that exhibited polar and/or side labelling (purple), purely polar labelling (blue), or a random labelling pattern (grey) as measured after 5 days of growth on YP agar. Wild type: n = 163 labelled hyphae; Δ*mreB1*: n = 184 labelled hyphae. **(D)** Representative DIC and epifluorescence (TRITC/Red) images of liquid YP-grown wild type strain labelled with the fluorescent TDL (47 ms exposure).

**Figure S6:**
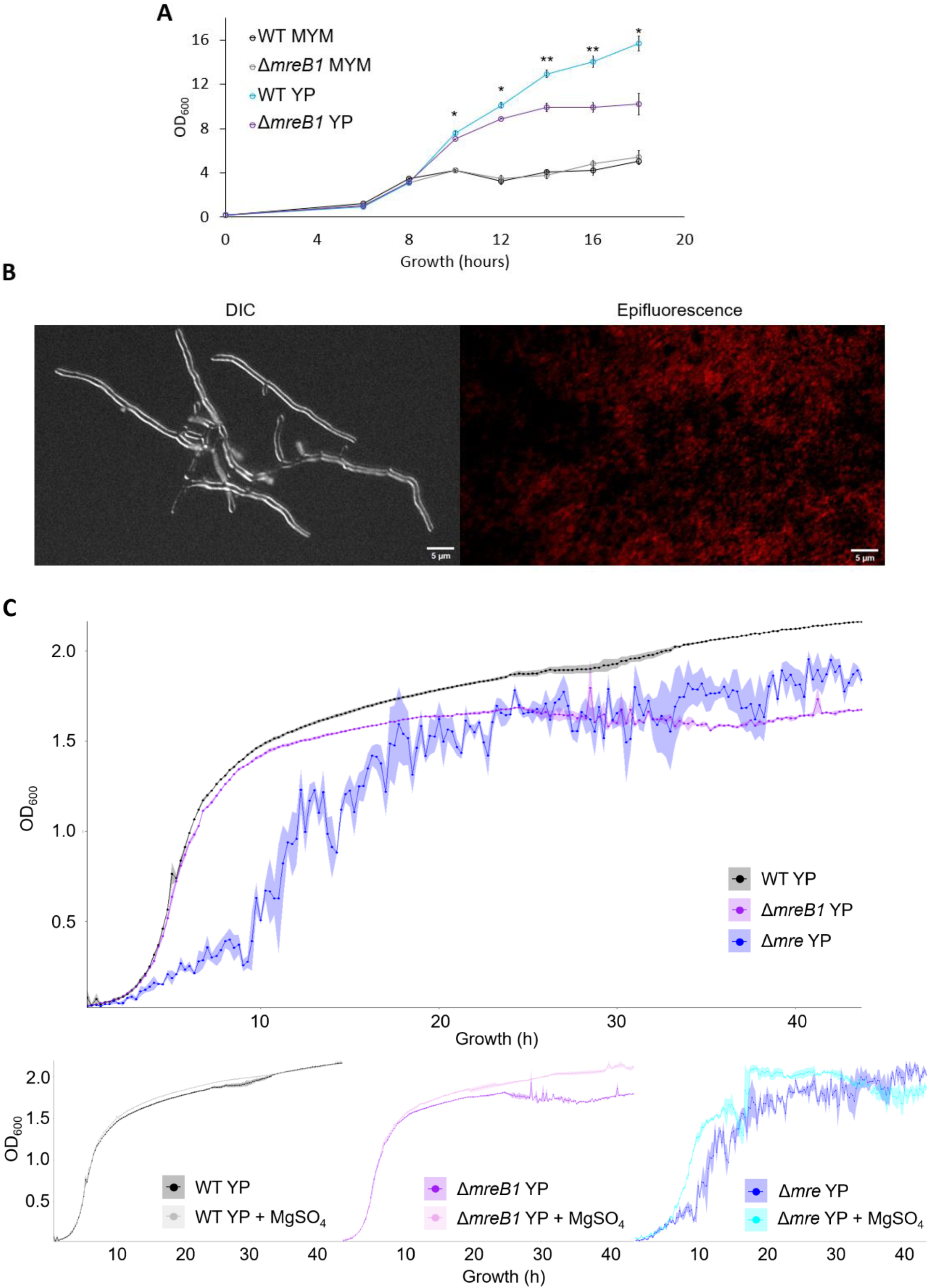
Enhanced *S. venezuelae* growth and *mreB* growth defects are observed in liquid YP medium and Mg rescues mutant defects in liquid YP. **(A)** OD_600_ growth curve comparing growth of the wild type (WT) and Δ*mreB1* strains in liquid MYM (classical growth) and YP (exploration-like) growth media. Cultures were sampled every two hours starting at hour 6 for a total of 18 hours of growth. Asterisks denote statistical significance between the wild type and Δ*mreB1* strains in liquid YP as determined by a Student’s t-test. No statistical differences were observed between wild type and Δ*mreB1* strains in classical (MYM) conditions. **(B)** Bottom right: wild type *S. venezuelae* (with no Halo tag being expressed) labelled with JaneliaFluor 549 as a negative control for autofluorescence and non-specific labelling with the HaloTag ligand (124 ms exposure). **(C)** Top: OD_600_ growth curves of the wild type, Δ*mreB1* and Δ*mre* strains in liquid YP. Bottom (left to right): OD_600_ growth curves of the wild type, Δ*mreB1* and Δ*mre* strains in liquid YP with and without supplementation with 25 mM MgSO_4_. For all curves, ribbon width represents standard error. Each point represents the average value of three independent biological replicates.

